# DNMT3B PWWP mutations cause hypermethylation of heterochromatin

**DOI:** 10.1101/2022.12.19.521050

**Authors:** Francesca Taglini, Ioannis Kafetzopoulos, Kamila Irena Musialik, Heng Yang Lee, Yujie Zhang, Mattia Marenda, Lyndsay Kerr, Hannah Finan, Cristina Rubio-Ramon, Hannah Wapenaar, Hazel Davidson-Smith, Jimi Wills, Laura C. Murphy, Ann Wheeler, Marcus D. Wilson, Duncan Sproul

## Abstract

The correct establishment of DNA methylation patterns is vital for mammalian development and is achieved largely by the *de novo* DNA methyltransferases DNMT3A and DNMT3B. Mutations in DNMT3B can cause immunodeficiency-centromeric instability-facial anomalies type 1 (ICF1) syndrome which is characterised by hypomethylated heterochromatin. However, in the genome, DNMT3B primarily localises to actively transcribing gene bodies through the interaction of its PWWP domain with the histone modification H3K36me3 and it is unclear how it is recruited to heterochromatin. Here we show that in DNMT3B knockout cells, loss of DNA methylation predominantly occurs in heterochromatic domains marked by H3K9me3. We also find that PWWP domain mutations which disrupt DNMT3B’s interaction with H3K36me3 result in striking increases of DNA methylation in H3K9me3-marked heterochromatin. Gains of methylation are also observed when the PWWP domain of DNMT3B is deleted. In contrast, we find that the ICF1 syndrome-causing PWWP mutation, S270P, does not result in hypermethylation of heterochromatin and destabilises the protein. We also show that removal of the N-terminus region of DNMT3B affects its recruitment to chromatin and ability to methylate H3K9me3 marked regions. Our results suggest that DNMT3B is recruited to H3K9me3 marked heterochromatin in a PWWP-independent manner and that this recruitment is facilitated by the protein’s N-terminus. More generally, we suggest that DNMT3B plays a role in DNA methylation homeostasis at heterochromatin, a process which is disrupted in ICF syndrome, cancer and aging.

## Introduction

DNA methylation is a repressive epigenetic mark that occurs predominantly on the cytosines of CpG dinucleotides in mammals. The mammalian genome is heavily methylated with the exception of some regulatory elements, particularly CpG islands (Suzuki and Bird, 2008). Studies in mouse showed that the DNA methylation landscape is established during early development by the *de novo* DNA methyltransferases (DNMTs), DNMT3A and DNMT3B (Okano et al., 1999). This developmentally established pattern is then largely maintained by the maintenance DNA methyltransferase DNMT1 (Goll and Bestor, 2005). However, some methylation maintenance is performed by the *de novo* DNMTs (Elliott et al., 2016; Liang et al., 2002; Liao et al., 2015).

In addition to their catalytic domains, DNMT3A and DNMT3B each possess chromatin reading ADD (ATRX-Dnmt3-Dnmt3L) and PWWP (Pro-Trp-Trp-Pro) domains which mediate their recruitment to the genome through interaction with histone modifications (Jeltsch and Jurkowska, 2016). The ADD domain of DNMT3A interacts with unmodified H3K4 (Otani et al., 2009; Zhang et al., 2010) to allosterically activate the protein (Guo et al., 2015; Li et al., 2011). This antagonizes DNMT3A’s activity at H3K4me3-marked promoters (Hu et al., 2009). The PWWP domains of both proteins bind to methylated H3K36 (Dhayalan et al., 2010; Morselli et al., 2015; Rondelet et al., 2016). DNMT3A’s PWWP domain preferentially binds H3K36me2, found at intergenic regions (Weinberg et al., 2019) whereas DNMT3B’s PWWP domain shows preference for H3K36me3 (Rondelet et al., 2016; Weinberg et al., 2019), recruiting the protein to the body of actively transcribed genes (Baubec et al., 2015; Morselli et al., 2015; Neri et al., 2017).

*Dnmt3b* is essential for development and knockout mice die at E11.5 (Okano et al., 1999). In humans, *DNMT3B* is mutated in the recessive Mendelian disorder immunodeficiency centromeric instability and facial abnormalities type 1 syndrome (ICF1) (Okano et al., 1999; Xu et al., 1999). The majority of missense mutations causing ICF1 syndrome occur in the catalytic domain of DNMT3B (Weemaes et al., 2013). A mutation of the PWWP domain, S270P, has also been identified in ICF1 (Shirohzu et al., 2002). ICF1 is characterised by widespread hypomethylation of satellite repeats and other heterochromatic loci (Heyn et al., 2012; Jeanpierre et al., 1993) and Dnmt3b localises to heterochromatic chromocenters by microscopy in mouse cells (Bachman et al., 2001). These constitutive heterochromatic loci are associated with the histone modification H3K9me3 (Allshire and Madhani, 2018; Brandle et al., 2022).

The majority of Dnmt3b localises to H3K36me3 by ChIP-seq (Baubec et al., 2015) and it is unclear how the protein is recruited to H3K9me3-marked constitutive heterochromatin. However, removal of the PWWP domain is reported to abolish DNMT3B localisation to chromocenters in mouse cells (Chen et al., 2004; Ge et al., 2004). The ICF1 syndrome S270P mutation has also been suggested to impair localisation to chromocenters (Chen et al., 2004; Ge et al., 2004) suggesting that the PWWP domain may play a role in DNMT3B’s recruitment to heterochromatin in addition to its better described role at H3K36me3.

To understand how DNMT3B is recruited to the genome, we have analysed *DNMT3B* knockout cells alongside DNMT3B mutations. We show that loss of *DNMT3B* disproportionality results in hypomethylation of heterochromatic H3K9me3 domains, and that mutations that disrupt PWWP-mediated H3K36me3 or DNA binding cause increased DNA methylation at H3K9me3-marked heterochromatin. Our results suggest that DNMT3B is recruited to constitutive H3K9me3-marked heterochromatin in a PWWP-independent manner that is facilitated by its N-terminus.

## Results

### DNMT3B methylates heterochromatin

To understand the role of DNMT3B in methylating different parts of the genome we examined DNMT3B knockout HCT116 cells (DNMT3B KO) (Rhee et al., 2002) using whole genome bisulfite sequencing (WGBS). Overall methylation levels were decreased to 96% and 86% of the levels observed in HCT116 wild-type cells by WGBS and mass spectrometry respectively (*Fig. S1a*).

To determine which type of genomic region was most affected by loss of DNMT3B we performed ChIP-seq for the histone modifications: H3K4me3 (promoter-associated), H3K36me3 (gene-body associated) and H3K9me3 (constitutive heterochromatin associated) in HCT116 cells. We also profiled the facultative heterochromatin-associated histone mark H3K27me3 which is deposited by the polycomb repressive complex 2 (Blackledge et al., 2015). We then correlated the levels of these histone modifications to changes in DNA methylation between HCT116 and DNMT3B KO cells in 2.5 kb windows across the genome (*Fig. 1a, S1b*). Losses of DNA methylation in DNMT3B KO compared to HCT116 were significantly correlated with each of the modifications (*Fig. S1b*, Pearson correlations, all p < 2.2×10^−16^). However, the correlations were much stronger with the constitutive H3K9me3 and facultative H3K27me3 heterochromatin modifications (Pearson correlations, R=-0.124 and -0.173 respectively, *Fig. S1b*) than with H3K36me3 and H3K4me3 (Pearson correlations, R=-0.053 and -0.030 respectively, *Fig. S1b*).

**Figure 1.**
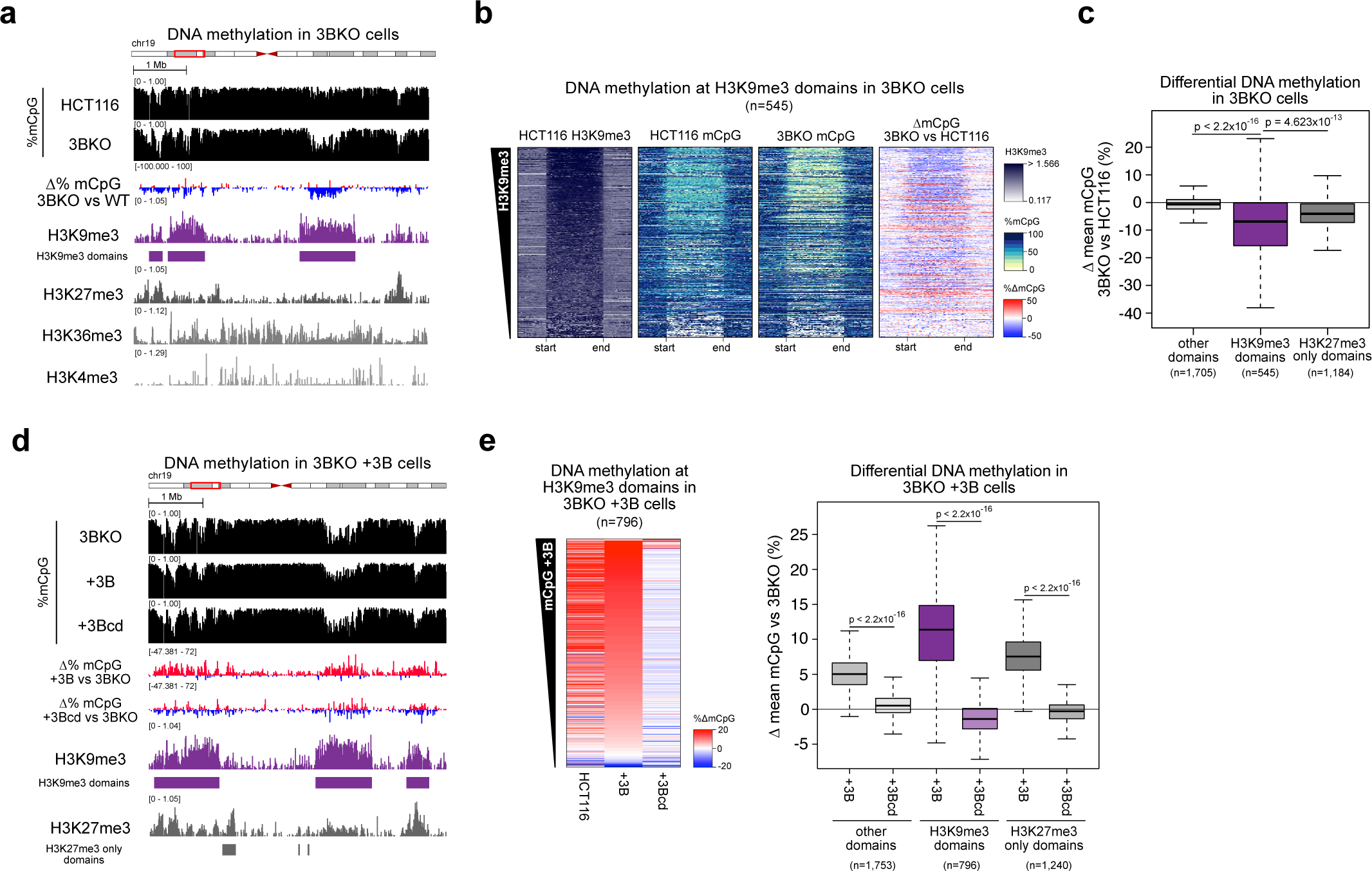
DNMT3B methylates heterochromatin. **a-c)** Loss of DNA methylation in DNMT3B KO cells occurs predominantly at heterochromatic domains. **a)** Representative genomic location showing loss of DNA methylation at H3K9me3-marked domains in DNMT3B KO cells (3BKO). Genome browser plots showing absolute (black) and differential (gain=red/loss=blue) DNA methylation levels alongside ChIP-seq and for histone modifications from HCT116 cells. ChIP-seq are normalised reads per 10^6^. H3K9me3 domains are defined in HCT116 cells. **b)** Heatmaps showing levels of H3K9me3 and of absolute or differential DNA methylation at HCT116 H3K9me3 domains, in HCT116 and DNMT3B KO cells. The domains are ranked by their mean H3K9me3 levels. **c)** Boxplot showing differential DNA methylation between DNMT3B KO and HCT116 at H3K9me3, H3K27me3-only domains and the domains marked by neither modification (other domains) defined in HCT116 cells. **d,e)** H3K9me3 domains gain DNA methylation when DNMT3B is re-expressed in DNMT3B KO cells. **d)** Representative genomic location showing gain of DNA methylation at H3K9me3 domains in DNMT3B KO cells expressing DNMT3B (+3B) or DNMT3B catalytically dead (+3Bcd). Genome browser plots show DNA methylation levels (absolute in black and gain=red/loss=blue), DNMT3B KO ChIP-seq signals and domains defined in DNMT3B KO cells. ChIP-seq are normalised reads per 10^6^. **e)** Left, heatmaps of relative DNA methylation levels at H3K9me3 domains. Values denote the change in DNA methylation relative to DNMT3B KO cells; H3K9me3 domains are defined in DNMT3B KO cells and ranked by the mean gain of methylation in DNMT3B KO cells expressing DNMT3B. Right, boxplots of DNA methylation difference to DNMT3B KO cells at H3K9me3, H3K27me3-only marked domains and domains not marked by either modification (other domains). For boxplots: Lines = median; box = 25th–75th percentile; whiskers = 1.5 × interquartile range from box. P-values are from two-sided Wilcoxon rank sum tests.

Given previous observations that DNMT3B is recruited to H3K36me3 (Baubec et al., 2015), we next examined gene bodies which are the primary locations marked by H3K36me3 (Bannister et al., 2005). Despite the role of DNMT3B at H3K36me3, the correlation between DNMT3B KO DNA methylation loss and H3K36me3 levels in gene bodies was very low and non-significant (Pearson correlation, R=-0.003, p-value = 0.671, *Fig. S1c*).

We then further examined the relationship between DNMT3B loss and the heterochromatin-associated H3K9me3 and H3K27me3 histone modifications. H3K9me3 marked broad genomic domains of reduced DNA methylation in HCT116 cells (*Fig. 1a*). In support of this, we found that hidden Markov model defined H3K9me3 domains significantly overlapped with extended domains of overall reduced methylation termed partially methylated domains (PMDs) defined in our HCT116 WGBS (Jaccard=0.575, p=1.07×10^-6^, Fisher’s test). They also significantly overlapped heterochromatic regions resistant to nuclease digestion identified using Protect-seq in the same cells (Jaccard=0.623, p=2.11×10^-30^, Fisher’s test) (Spracklin and Pradhan, 2020). HCT116 H3K9me3 domains lost significantly more methylation in DNMT3B KO cells than non-heterochromatic domains which were marked by neither H3K9me3 nor H3K27me3 (other domains, p < 2.2×10^-16^ Wilcoxon test, *Fig. 1a, b and c*). Although HCT116 H3K9me3 domains showed some enrichment for H3K27me3 (*Fig. S1d*), facultative heterochromatic domains marked by H3K27me3 alone lost DNA methylation to a significantly lesser degree in DNMT3B KO cells than H3K9me3 domains (p < 1.8×10^-9^ Wilcoxon test, *Fig. 1c*). This suggests that DNMT3B is more active in regions of constitutive heterochromatin marked by H3K9me3 than in facultative heterochromatin marked by H3K27me3.

To confirm that DNMT3B methylates DNA at H3K9me3-marked heterochromatin we expressed the major catalytically active DNMT3B isoform expressed in somatic cells, DNMT3B2 (Weisenberger et al., 2004), in DNMT3B KO cells to generate DNMT3B^WT^ cells (DNMT3B2 is henceforth referred to as DNMT3B, *Fig. S1e*, top). We then compared gains of DNA methylation to histone modification levels assayed by ChIP-seq from DNMT3B KO cells. H3K9me3 domains were also observed in DNMT3B KO cells, and we defined them using a hidden Markov model (*Fig. 1d*). DNMT3B KO H3K9me3 domains significantly overlapped HCT116 H3K9me3 domains (Jaccard=0.582, p=9.82×10^-162^, Fisher’s exact test) but showed a greater enrichment of H3K27me3 than HCT116 H3K9me3 domains (*Fig. S1d* and *f*). Changes in DNA methylation in DNMT3B^WT^ cells significantly correlated with H3K27me3 and H3K9me3 (Pearson correlations, R=0.306 and 0.230 respectively, both p < 2.2×10^−16^) and more weakly with H3K36me3 and H3K4me3 (Pearson correlations, R=0.027 and 0.143 respectively, both p < 2.2×10^−16^). In DNMT3B^WT^ cells, gains of DNA methylation at DNMT3B KO H3K9me3 domains were significantly greater than those in domains defined by H3K27me3 alone or lacking either mark (*Fig. 1d, e*), consistent with greater DNMT3B activity at H3K9me3 than H3K27me3 marked heterochromatin. Gains of DNA methylation in DNMT3B^WT^ cells were also significantly greater than those observed upon expression of a catalytically inactive mutant DNMT3B (Hsieh, 1999) (*Fig. 1d, e*).

To determine whether activity at H3K9me3 was specific to DNMT3B, we also expressed DNMT3A1, the DNMT3A isoform expressed in somatic cells (Chen et al., 2002), and catalytically inactive DNMT3A1 (Hsieh, 1999), in DNMT3B KO cells (*Fig. S1e*, bottom). The DNA methylation gains observed upon expression of DNMT3B were significantly greater at H3K9me3 domains compared to the DNA methylation gains induced by DNMT3A and catalytically inactive DNMT3A1 (*Fig. S1g, h*). In contrast, the weaker gains of DNA methylation observed at H3K27me3 domains were more similar between DNMT3B and DNMT3A (*Fig. S1g, h*).

Overall, these data from the knockout and re-expression of DNMT3B suggest that DNMT3B specifically methylates constitutive heterochromatic domains marked by H3K9me3.

### Disruption of DNMT3B H3K36me3 binding causes gains of DNA methylation at heterochromatin

Having observed a role for DNMT3B in methylating constitutive H3K9me3-marked heterochromatin throughout the genome, we sought to understand the mechanism(s) responsible for its recruitment to this genomic compartment. Previous work has suggested a role for DNMT3B’s PWWP in localising it to constitutive heterochromatic chromocenters by microscopy (Chen et al., 2004; Ge et al., 2004). To impair the key H3K36me3 binding function of DNMT3B’s PWWP we therefore mutated a key residue of its aromatic cage, tryptophan 263 to alanine (W263A) (*Fig. 2a*) and expressed this protein in DNMT3B KO cells (DNMT3B^W263A^ cells). Mutation of the paralogous residue in DNMT3A disrupts H3K36me2/3 binding and causes Heyn-Sproul-Jackson Syndrome (HESJAS) and (Heyn et al., 2019), and DNMT3B tryptophan 263 contacts the methylated H3 lysine 36 (*Fig. S2a*).

**Figure 2.**
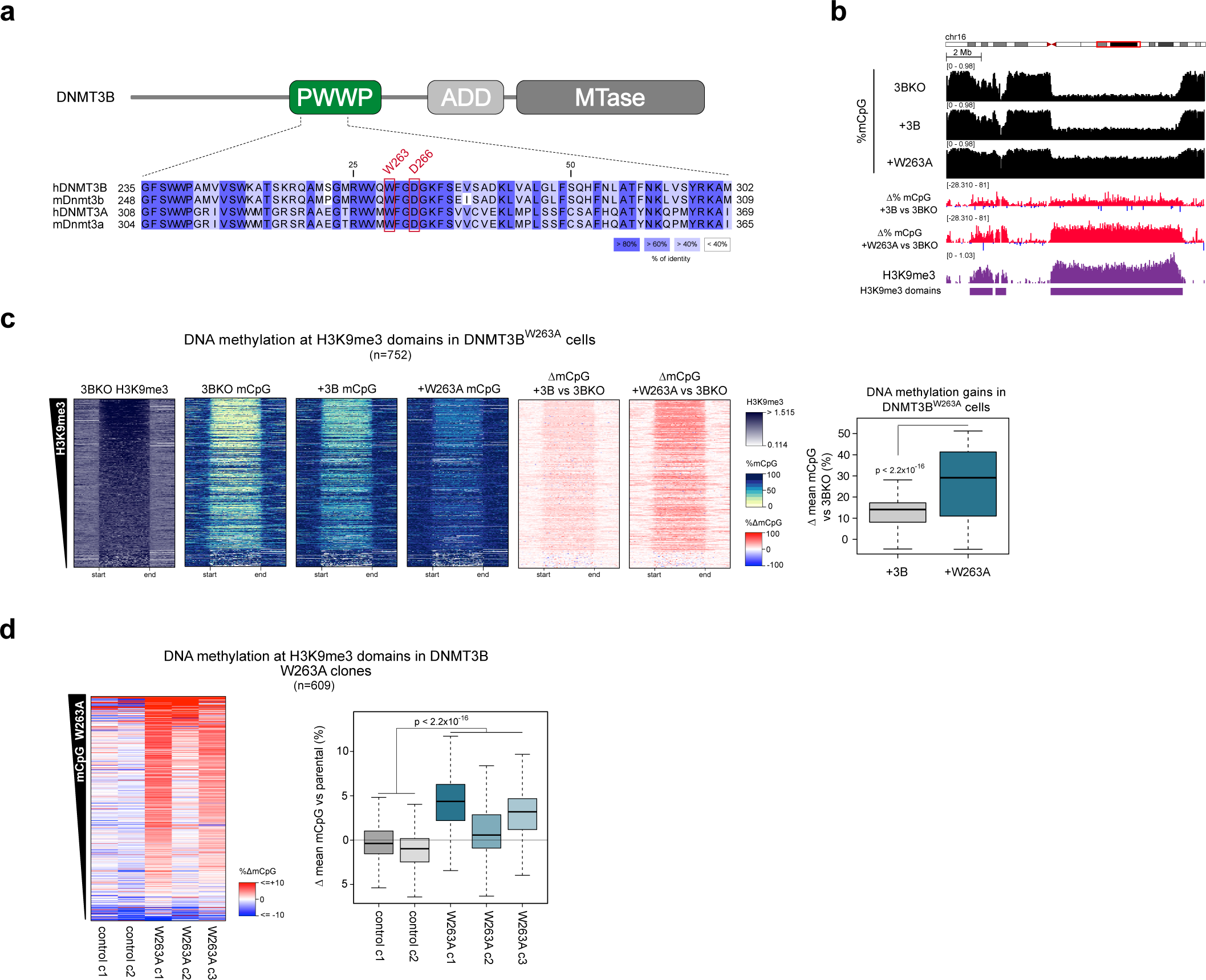
Disruption of DNMT3B H3K36me3 binding causes gains of DNA methylation at heterochromatin. **a)** Schematic of DNMT3B protein and domains (top) and part of a multiple sequence alignment of DNMT3B’s PWWP domain with the DNMT3A and mouse orthologues (bottom). **b,c)** DNMT3B^W263A^ remethylates heterochromatin to a significantly higher level than DNMT3B^WT^. **b)** Genome browser plots showing absolute (black) and differential (gain=red/loss=blue) DNA methylation levels, DNMT3B KO ChIP-seq signals at H3K9me3 domains in DNMT3B KO, DNMT3B^WT^ and DNMT3B^W263A^ cells. ChIP-seq are normalised reads per 10^6^. **c)** Left, heatmaps showing levels of H3K9me3 and DNA methylation at H3K9me3 domains in DNMT3B KO, DNMT3B^WT^ and DNMT3B^W263A^ cells. The domains are ranked by their mean H3K9me3 levels. Right, boxplot showing gains of DNA methylation at H3K9me3 domains in DNMT3B^WT^ and DNMT3B^W263A^ cells. **d)** Endogenous expression of DNMT3B^W263A^ in HCT116 cells leads to hypermethylation of heterochromatin. Left, heatmaps of DNA methylation levels at H3K9me3 domains in CRISPR/Cas9-edited DNMT3B^W263A^ and control clones relative to parental HCT116 cells. H3K9me3 domains are defined in HCT116 cells and ranked by the mean gain of DNA methylation in DNMT3B^W263A^ clones. Right, boxplot showing differential DNA methylation at H3K9me3 domains in CRISPR clones compared to the parental cell line. For all the boxplots: lines = median; box = 25th–75th percentile; whiskers = 1.5 × interquartile range from box. P-values are from two-sided Wilcoxon rank sum tests.

DNMT3B^W263A^ was expressed to a lower level than DNMT3B^WT^ in DNMT3B KO cells (*Fig. S2b*). We therefore checked for protein stability using a biscistronic reporter expressing GFP-DNMT3B^WT^ or GFP-DNMT3B^W263A^ and dsRed (Huang et al., 2022) in DNMT3B KO cells. In this system GFP intensity reports DNMT3B levels and transfection efficiency is reported by dsRed intensity. The mean ratio of GFP to dsRed intensity in DNMT3B^W263A^ cells was 91.6% of that observed in DNMT3B^WT^ cells (*Fig. S2c*), indicating that the proteins are similarly stable. In addition, the melting temperature of DNMT3B^W263A^ was similar to that of DNMT3B^WT^ *in vitro* when constructs consisting of the PWWP and ADD domain were assayed by thermal denaturation assays (46.7 and 49.9°C respectively, *Fig. S2d*).

We then assessed the effect of DNMT3B^W263A^ on the methylome using WGBS. Despite the lower protein level observed, global levels of methylation were higher in DNMT3B^W236A^ cells than in DNMT3B^WT^ cells (*Fig. S2e*). Surprisingly given the PWWP domains previous association with heterochromatic localisation (Chen et al., 2004; Ge et al., 2004), gains of DNA methylation in DNMT3B^W263A^ cells versus DNMT3B KO cells were highly significantly correlated with H3K9me3 levels in DNMT3B KO cells (Pearson correlation, R=0.516, p < 2.2×10^−16^, *Fig. S2f*, right). This correlation and the absolute gains of DNA methylation were greater than differences between DNMT3B^WT^ and DNMT3B KO cells (Pearson correlation, R=0.299, p < 2.2×10^-16^, *Fig. S2f*, left). In support of this, significantly more methylation was present at heterochromatic H3K9me3 domains in DNMT3B^W263A^ cells than DNMT3B^WT^ cells (*Fig. 2b* and *c*).

Having seen gains of methylation at interspersed heterochromatin in DNMT3B^W326A^ cells, we then asked if similar gains were seen at constitutively heterochromatic satellite II repeats which are hypomethylated in ICF1 syndrome (Hassan et al., 2001; Xu et al., 1999). Using a methylation-sensitive Southern blot, we observed greater digestion of satellite II repeats in DNMT3B KO cells as compared to HCT116 cells, consistent with hypomethylation of satellite II repeats upon loss of DNMT3B (*Fig. S2g*). Protection from digestion was restored by expression of DNMT3B^WT^ (*Fig. S2g*). However, DNMT3B^W263A^ cells displayed an even greater protection from digestion indicating that the mutation results in hypermethylation of satellite II compared to DNMT3B^WT^ (*Fig. S2g*).

To validate that the hypermethylation of heterochromatin in DNMT3B^W263A^ cells was caused by the disruption H3K36me3 binding, we then mutated another residue of the DNMT3B PWWP’s aromatic cage, D266, to alanine (*Fig. 2a* and *S2a*) (Dhayalan et al., 2010), and expressed this in DNMT3B KO cells (DNMT3B^D266A^ cells, *Fig. S2b*). By methylation-sensitive Southern blot, satellite II repeats were similarly protected from digestion in DNMT3B^D266A^ cells as in DNMT3B^W263A^ cells (*Fig. S2g*). We also assessed the methylation level of two representative non-repetitive loci within heterochromatic H3K9me3 domains (*Fig. S2h*) by bisulfite PCR, finding them to be significantly more methylated in DNMT3B^W263A^ and DNMT3B^D266A^ cells than in DNMT3B^WT^ cells or DNMT3B KO cells expressing GFP (*Fig. S2i*).

To confirm that the hypermethylation of H3K9me3-marked heterochromatin observed in DNMT3B^W263A^ cells was not a result of ectopically elevated protein levels, we generated 3 homozygous *DNMT3B^W263A^* knock-in clonal lines from HCT116 cells using CRISPR-Cas9 (*Fig. S2j*). We then compared their DNA methylation pattern to two similarly treated control cell line clones which lacked the mutation using WGBS. DNMT3B^W263A^ knock-in clones showed gains of DNA methylation at H3K9me3 domains compared to parental HCT116 cells, and these gains were significantly greater than the small differences in DNA methylation seen in control clones (*Fig. 2d*).

These results therefore suggest that disruption of DNMT3B’s H3K36me3 binding through the PWWP domain results in hypermethylation of H3K9me3-marked heterochromatin.

### Impaired H3K36me3 binding leads to increased DNMT3B localisation at heterochromatin

To understand the localisation of DNMT3B^W263A^ in more detail, we performed ChIP-seq analysis of DNMT3B^W263A^ cells and compared this to DNMT3B^WT^ cells.

We first checked that the ectopically expressed T7-tagged DNMT3B localised similarly to endogenous DNMT3B by comparing ectopic DNMT3B localisation in HCT116 cells to our previously published endogenously tagged ChIP-seq of DNMT3B in these cells (Masalmeh et al., 2021). ChIP-seq signal for ectopically expressed DNMT3B was significantly correlated with that of endogenously tagged DNMT3B (Pearson correlation, R=0.630, p < 2.2×10^−16^, *Fig. S3a* and *b*). ChIP-seq signal for both also correlated significantly with H3K36me3 signal across the genome (Pearson correlations R=0.395 and 0.550 respectively, both p < 2.2×10^−16^, *Fig. S3a*) suggesting that ectopically expressed DNMT3B recapitulates the localisation of the endogenous protein.

We then examined the effect of the W263A mutation on DNMT3B localisation in DNMT3B KO cells. The ChIP-seq signal from DNMT3B^WT^ cells correlated with levels of DNMT3B KO H3K36me3 across the genome (Pearson correlation, R=0.386, p < 2.2×10^−16^, *Fig. S3c* and *3a*) and in gene bodies (*Fig. S3d*). In contrast, levels at gene bodies in DNMT3B^W263A^ cells were far lower than DNMT3B^WT^ cells (*Fig. 3a* and *S3d*). However, DNMT3B^W263A^ was seen at some H3K36me3-marked loci (*Fig. 3a*, orange arrows) and low enrichment at gene bodies marked by the highest levels of H3K36me3 could still be detected (*Fig. S3d*). This suggests DNMT3B^W263A^ impairs but does not completely abolish localisation to H3K36me3.

**Figure 3.**
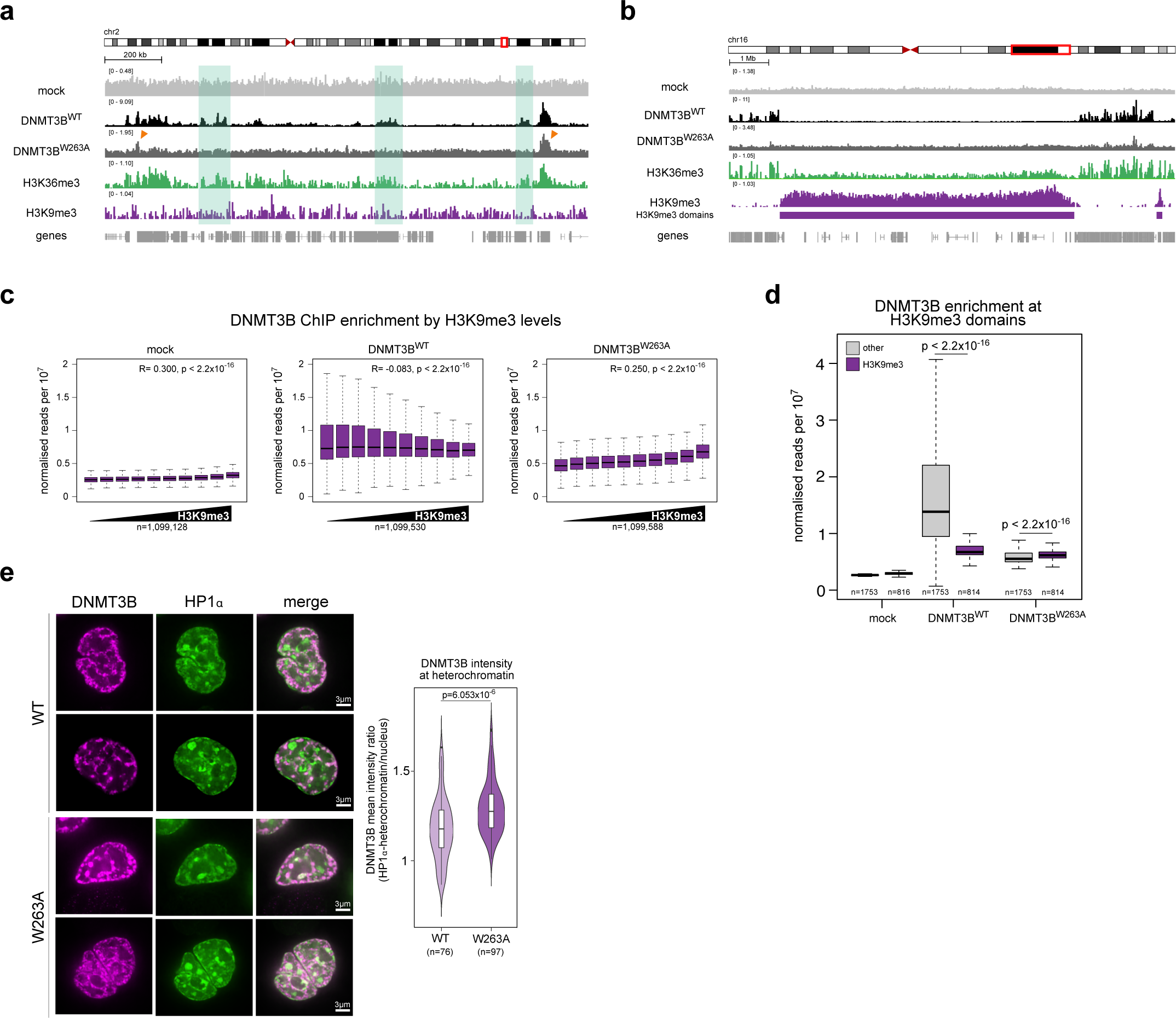
Impaired H3K36me3 binding leads to increased DNMT3B localisation to heterochromatin. **a)** DNMT3B^WT^ and DNMT3B^W263A^ binding profiles at a representative euchromatic genomic region. **b)** DNMT3B^WT^ and DNMT3B^W263A^ binding profiles at a representative heterochromatic genomic region. Both **a.** and **b.** are genome browser plots showing T7-DNMT3B ChIP signal along with H3K36me3 and H3K9me3 ChIP-seq in DNMT3B KO cells, normalised reads per 10^6^. In **a.** green rectangles highlight DNMT3B^W263A^ loss from several H3K36me3-marked regions; orange arrows indicate remaining DNMT3B^W263A^ peaks at some H3K36me3 loci. **c,d)** DNMT3B^W263A^ has increased localisation at H3K9me3 domains. **c)** T7 ChIP-seq counts at 2.5 kb genomic windows were ranked by increased H3K9me3 levels measured in DNMT3B KO cells before being divided into deciles. Pearson’s correlations (R) and associated p-values are shown. **d)** Boxplot showing T7-DNMT3B enrichment at H3K9me3 domains and genomic regions marked by neither H3K9me3 nor H3K27me3 (other). **e)** DNMT3B^W263A^ localises to HP1⍺. Right, representative confocal images of SNAP-DNMT3B and HP1⍺-GFP localisation in live DNMT3B KO cells. Bars = 3 µm. Left, violin plot showing the distribution of DNMT3B mean intensity ratio between HP1⍺-marked heterochromatin and the rest of the nucleus from one representative experiment. For the boxplots: lines = median; box = 25th–75th percentile; whiskers = 1.5 × interquartile range from box. Boxplot p-values are from two-sided Wilcoxon rank sum tests.

In contrast to its correlation with H3K36me3, DNMT3B^WT^ levels negatively correlated with DNMT3B KO H3K9me3 levels genome-wide (Pearson’s correlation, R=-0.083, p < 2.2×10^−16^, *Fig. 3b* and **c**). We did not detect obvious peaks of DNMT3B^W263A^ enrichment within H3K9me3 domains (*Fig. 3b*). However, in contrast to DNMT3B^WT^, DNMT3B^W263A^ signal was positively correlated with H3K9me3 signal (Pearson’s correlation, R=0.250, p < 2.2×10^−16^, *Fig. 3c*). In agreement with this, DNMT3B^W263A^ enrichment at H3K9me3 domains was significantly higher than at other domains, whereas DNMT3B^WT^ enrichment was significantly lower at H3K9me3 domains than at other domains (*Fig. 3d*). This is consistent with increased DNMT3B^W263A^ localisation at H3K9me3 and our observations of increased DNA methylation at H3K9me3 in these same cells.

To further understand the localisation of DNMT3B^W263A^ on chromatin using an independent approach, we compared the localisation of SNAP-tagged DNMT3B^WT^ and DNMT3B^W263A^ in live DNMT3B KO cells using confocal microscopy. Both proteins were broadly distributed in the nucleus and showed variable patterns (*Fig. 3e*). To assay their degree of localisation at heterochromatin, we then compared their distribution to a marker of H3K9me3-marked constitutive heterochromatin, HP1*α*-GFP (*Fig. 3e*). The ratio of DNMT3B signal at HP1*α* -marked heterochromatin versus the rest of nucleus was significantly higher for DNMT3B^W263A^ than for DNMT3B^WT^, consistent with a greater localisation of DNMT3B^W263A^ to H3K9me3-marked heterochromatin (*Fig. 3e* and *S3e*). We also assessed the degree of localisation at another constitutive heterochromatic compartment, the nuclear periphery (van Steensel and Belmont, 2017). Consistent with our ChIP-seq results, a significantly greater proportion of DNMT3B^W263A^ signal was found at the nuclear periphery than DNMT3B^WT^ signal (*Fig. S3f*).

Taken together these results suggest that interfering with the interaction of DNMT3B’s PWWP with H3K36me3 causes increased localisation of DNMT3B at H3K9me3-marked constitutive heterochromatin.

### The ICF1-associated PWWP S270P mutation destabilises DNMT3B

Having observed that mutations interfering with DNMT3B’s interaction with H3K36me3 resulted in increased localisation and gains of methylation at heterochromatin, we next sought to understand the effect of the ICF1-associated S270P mutation (Shirohzu et al., 2002) (*Fig. S4a*).

Similarly to the mutations we have examined, S270P has been reported to reduce the interaction of DNMT3B with H3K36me3-marked nucleosomes (Baubec et al., 2015). S270 is located distal to the methyl-lysine recognising binding pocket (*Fig. S4b*) and has been hypothesised to make a hydrogen bond with the backbone of bound H3 peptide (Rondelet et al., 2016). However, in contrast to our observations with other PWWP mutations, S270P has been reported to reduce pericentromeric localisation in mouse cells (Chen et al., 2004; Ge et al., 2004).

We therefore expressed DNMT3B^S270P^ in DNMT3B KO cells (DNMT3B^S270P^ cells) and asked whether it increased methylation at satellite II repeats by methylation sensitive Southern blot. In contrast to the increased protection from digestion observed in DNMT3B^WT^ and DNMT3B^W263A^ cells, the level of digestion in DNMT3B^S270P^ cells was similar to that of DNMT3B KO cells (*Fig. 4a*), suggesting that expression of DNMT3B^S270P^ did not lead to an increase of DNA methylation at satellite II. The levels of DNA methylation at two non-repetitive H3K9me3 loci in DNMT3B^S270P^ cells were also similar to those seen in DNMT3B KO cells expressing GFP when measured by bisulfite PCR (*Fig. 4b*).

**Figure 4.**
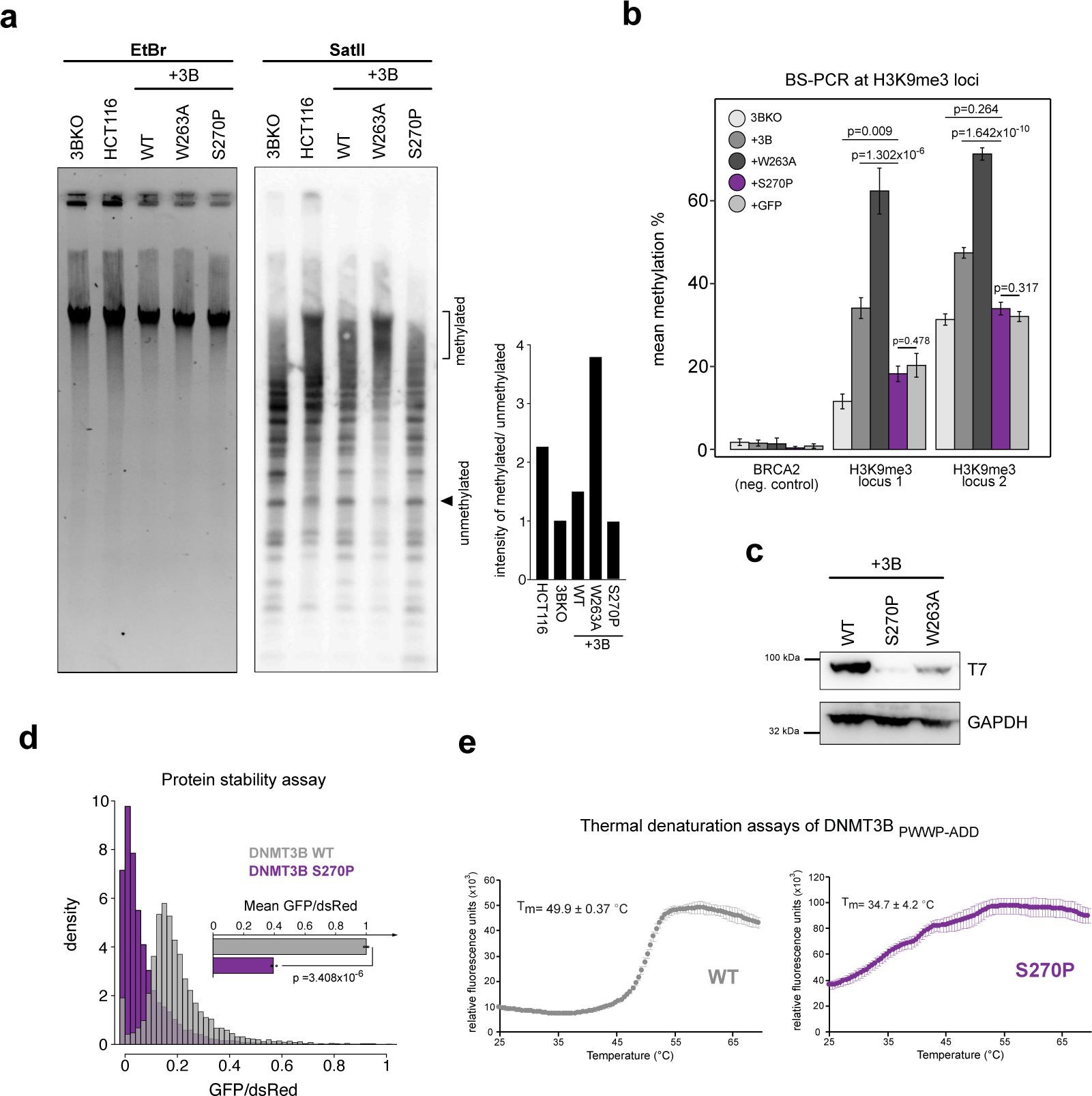
The ICF syndrome S270P mutation destabilises DNMT3B. **a,b)** DNMT3B^S270P^ cannot remethylate heterochromatin. **a)** Methylation sensitive Southern blot showing digestion of satellite II sequences in DNMT3B KO cells expressing DNMT3B mutants (centre). Ethidium bromide stained gel (EtBr) is shown as a loading control (left). Barplot shows signal quantification of satellite II Southern blot using the ratio of the methylated over unmethylated regions indicated (right). **b)** Mean methylation by BS-PCR at H3K9me3 loci alongside the H3K4me3-marked BRCA2 promoter in DNMT3B mutant cells. P-values are from two-sided Wilcoxon rank sum tests. The number of reads analysed per sample is shown in *Supplementary Table 1*. **c-e)** DNMT3B protein carrying S270P mutation is unstable. **c)** Representative western blot showing expression of DNMT3B proteins in DNMT3B KO cells. **d)** Stability of DNMT3B^WT^ and DNMT3B^S270P^ measured by fluorescence reporter. Histogram shows the density distribution of single cell GFP/dsRed ratios of one representative experiment. The barplot shows the mean GFP/dsRed ratios of all the cells, measurements for the mutant normalized to the wild-type, for 3 independent experiments. P value is from Student’s T-test. **e)** Thermal denaturation assays on purified DNMT3B^WT^ or DNMT3B^S270P^ protein PWWP and ADD fragments. Graphs showing SYBR-safe fluorescence measured at temperatures from 25°C to 69.5°C. Mean and standard deviation of three experiments are plotted with mean Tm values stated.

While these data were consistent with previous findings that S270P impairs DNMT3B recruitment to heterochromatin, they were inconsistent with its proposed effect on H3K36me3 binding and our observations in DNMT3B^W263A^ and DNMT3B^D266A^ cells. We noticed that DNMT3B^S270P^ cells expressed much lower levels of protein than DNMT3B^WT^ cells by western blot suggesting that it might not be stable (*Fig. 4c*). Mutations to prolines can often prematurely terminate secondary structures and destabilise proteins and S270 lies within a *β*-sheet integral to the PWWP domain. We therefore tested the stability of the DNMT3B^S270P^ mutant. Using our fluorescent reporter protein stability assay, we observed that the ratio of GFP to dsRed intensity in DNMT3B^S270P^ cells was significantly lower than DNMT3B^WT^ cells consistent with destabilisation of DNMT3B by S270P (*Fig. 4d*). In addition, *in vitro* a purified S270P mutant construct containing the PWWP and ADD domains was far less stable, as evidenced by the lower melting temperature of DNMT3B^S270P^ compared to DNMT3B^WT^ when assessed by thermal denaturation assays (34.7 and 49.9°C respectively, *Fig. 4e* and *S4c*).

These results suggest that the failure of DNMT3B^S270P^ to methylate heterochromatin is caused by the destabilising effect of the mutation rather than its effect on H3K36me3 binding.

### DNMT3B’s PWWP domain is dispensable for DNMT3B localisation to heterochromatin

Having observed that DNMT3B PWWP mutations affecting H3K36me3 binding result in increased localisation and methylation at H3K9me3 domains, we then sought to understand which parts of the protein might be responsible for recruiting DNMT3B to heterochromatin.

As well as methylated H3K36, PWWP domains are reported to bind DNA through positively charged surface residues (Eidahl et al., 2013; van Nuland et al., 2013). DNMT3B has also been reported to directly bind DNA (Chen et al., 2004). We therefore sought to understand whether DNA binding mediated through DNMT3B’s PWWP might be responsible for its recruitment to heterochromatin.

Analysis of the structure of DNMT3B’s PWWP domain reveals two patches of positively charged residues distal to the H3K36me3 binding pocket that might be involved in DNA binding (*Fig. 5a*, see methods for details). These contain two highly conserved lysines, K276 and K294, and two less conserved positively charged residues, K251 and R252 (*Fig. 5a* and *S5a*). We tested the role of these residues in DNA binding by EMSA and found that mutating all 4 amino acids drastically reduced DNA binding (*Fig. 5b* and *Fig. S5a*). Charge swapping mutations of Lys-276 or Lys-294 alone are however sufficient to highly decrease binding to DNA (*Fig. 5b* and *S5a*).

**Figure 5.**
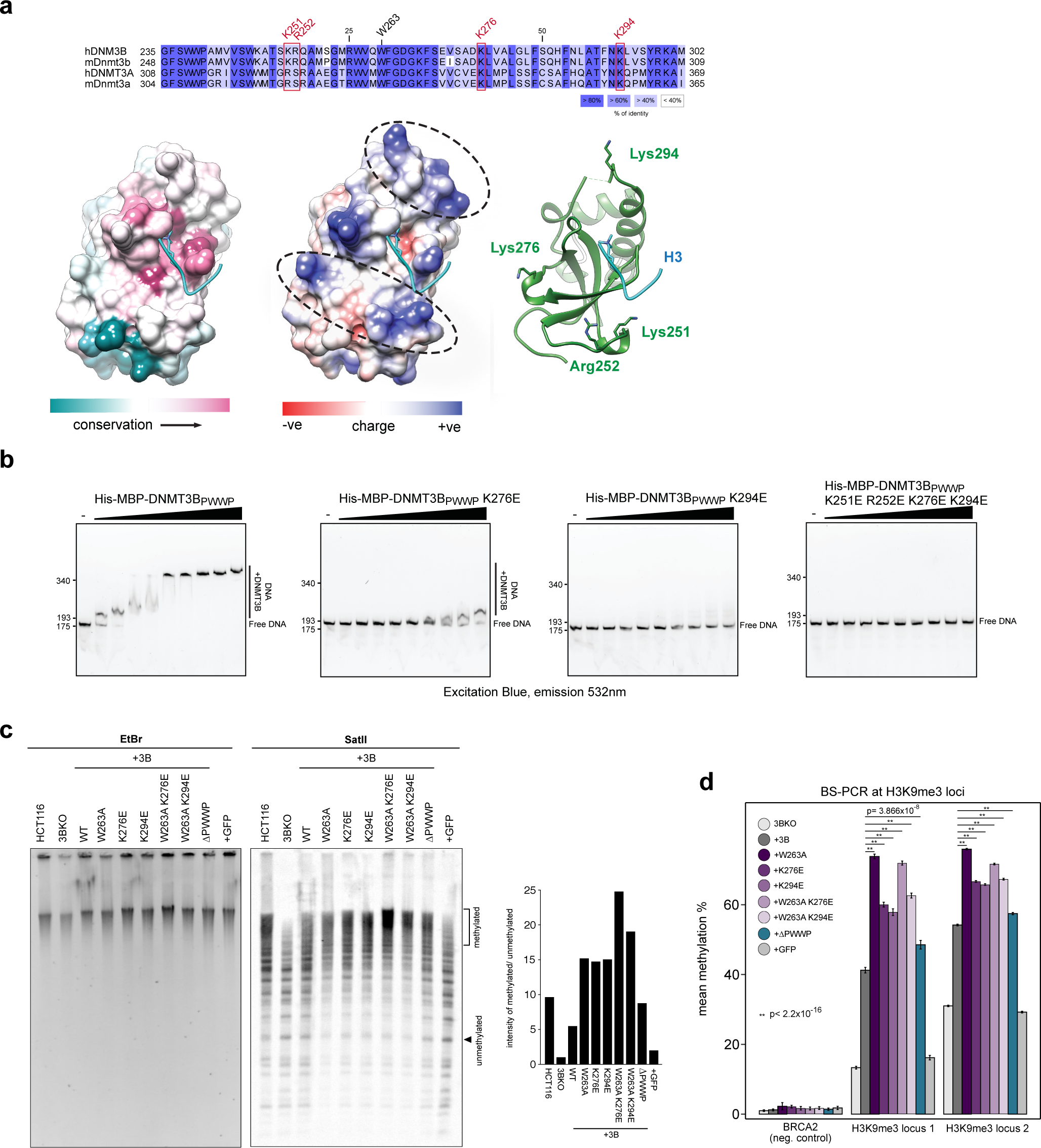
DNMT3B-PWWP binding to DNA is dispensable for localisation to heterochromatin. **a, b)** Positively charged residues on the surface of DNMT3B’s PWWP bind DNA. **a)** Sequence of the DNMT3B PWWP domain alongside its structure (PDB: 5CIU) (Rondelet et al., 2016). Surface representation coloured according to amino-acid sequence conservation calculated by ConSurf (left) (Ashkenazy et al., 2016), or columbic surface charge (middle). Dashed ovals indicate clusters of positive charge. Ribbon model of domain with putative DNA-binding basic residues mutated in this study highlighted (right). **b)** Representative gel images from electrophoretic mobility shift assays using increasing concentrations of tagged DNMT3B PWWP domains containing mutations with fluorescently labelled DNA. **c, d)** DNMT3B PWWP mutants affecting DNA binding remethylate heterochromatin to a greater extent than WT protein. **c)** Methylation sensitive Southern blot showing digestion of satellite II sequences in DNMT3B KO cells expressing DNMT3B mutants (centre). Ethidium bromide stained gel (EtBr) is shown as a loading control (left). Barplot shows signal quantification of satellite II Southern blot using the ratio of the methylated over unmethylated regions indicated (right). **d)** Mean methylation by BS-PCR at H3K9me3 loci alongside the H3K4me3-marked BRCA2 promoter in DNMT3B mutant cells. P-values are from two-sided Wilcoxon rank sum tests. The number of reads analysed per each sample are shown in *Supplementary Table 4*.

We therefore expressed these single mutations in DNMT3B KO cells (DNMT3B^K276E^ and DNMT3B^K294E^ cells respectively, *Fig. S5b*) and compared DNA methylation at satellite II sequences to DNMT3B^WT^ and DNMT3B^W263A^ cells by methylation sensitive Southern blot. Both DNMT3B^K276E^ and DNMT3B^K294E^ cells exhibited protection from digestion at a level similar to that observed for DNMT3B^W263A^ cells (*Fig. 5c*). Using bisulfite PCR we found that methylation levels at two non-repetitive H3K9me3 loci were significantly higher in DNMT3B^K276E^ or DNMT3B^K294E^ cells compared to DNMT3B^WT^ cells (*Fig. 5d*). These results suggest that these two mutations, that affect DNA binding, result in increased localisation of DNMT3B to heterochromatin similar to mutations affecting the interaction with H3K36me3. Expression of DNMT3B containing these lysine mutants alongside the W263A mutation (DNMT3B^W263A+K276E^ and DNMT3B^W263A+K276E^ cells) resulted in higher methylation levels than the single lysine mutations at both satellite II and H3K9me3 loci (*Fig. 5c* and *d*).

To further understand the requirement for the PWWP domain in localising DNMT3B to heterochromatin, we expressed DNMT3B lacking the entire PWWP domain (DNMT3B^ΔPWWP^). DNMT3B^ΔPWWP^ levels were lower than DNMT3B^WT^ by western blot (*Fig. S6a*). Similarly to DNMT3B^W263A^ cells, methylation sensitive Southern blot showed that in DNMT3B^ΔPWWP^ cells, satellite II was protected from digestion to a greater degree than in DNMT3B^WT^ cells (*Fig. 5c*). We also observed significantly greater levels of DNA methylation at two non-repetitive H3K9me3 loci in DNMT3B^ΔPWWP^ cells than in DNMT3B^WT^ cells (*Fig. 5d*). Although the degree of the hypermethylation in DNMT3B^ΔPWWP^ cells was less than observed for DNMT3B^W263A^ cells, this is consistent with the lower level DNMT3B^ΔPWWP^ compared to DNMT3B^W263A^.

Overall, these results suggest that interference with the DNA or H3K36me3 binding capacity of DNMT3B’s PWWP domain results in hypermethylation of heterochromatin and that the PWWP domain itself is dispensable for recruitment of DNMT3B to heterochromatin.

### The N-terminus facilitates methylation of heterochromatin by DNMT3B

Recently, the N-terminal region of DNMT3A has been shown to interact with H2AK119ub (Gu et al., 2022; Weinberg et al., 2021) and recruit the protein to H3K27me3-marked DNA methylation valleys, particularly in the context of PWWP domain mutations (Heyn et al., 2019; Manzo et al., 2017; Weinberg et al., 2021). We therefore wondered whether the N-terminal region of DNMT3B might be involved in its recruitment to heterochromatin, particularly given that this is the most divergent region between DNMT3A and DNMT3B (Manzo et al., 2017).

We expressed DNMT3B lacking the N-terminal region in DNMT3B KO cells (lacking residues 1 to 199, DNMT3B^ΔN^ cells, *Fig. S6a*) and assayed their methylation profile by WGBS (*Fig. 6a*). Methylation levels at H3K9me3 domains in DNMT3B^ΔN^ cells were significantly greater than in DNMT3B KO cells, but significantly lower than DNMT3B^WT^ cells in two independently generated cell lines (mean of two replicates in *Fig. 6a, b* and *Fig. S6b*). Levels of DNA methylation at other domains were not significantly different between DNMT3B^ΔN^ and DNMT3B^WT^ cells (*Fig. 6b*). Methylation sensitive Southern blot revealed that methylation levels at satellite II repeats in DNMT3B^ΔN^ cells were lower than in DNMT3B^WT^ cells, as seen by increased digestion due to higher levels of hypomethylated DNA (*Fig. S6c*). Significantly lower levels of DNA methylation were also observed at two non-repetitive H3K9me3 loci in DNMT3B^ΔN^ cells compared to DNMT3B^WT^ cells (*Fig. S6d*). Similarly, cells expressing a doubly mutated DNMT3B^W263A^ lacking the N-terminal region (DNMT3B^ΔN+W263A^ cells) showed less methylation at satellite II (*Fig. S6c*) and at the two selected non-repetitive H3K9me3 loci (*Fig. S6d*) compared to DNMT3B^W263A^ cells. These data suggest that the N-terminal region of DNMT3B facilitates its activity at heterochromatin.

**Figure 6.**
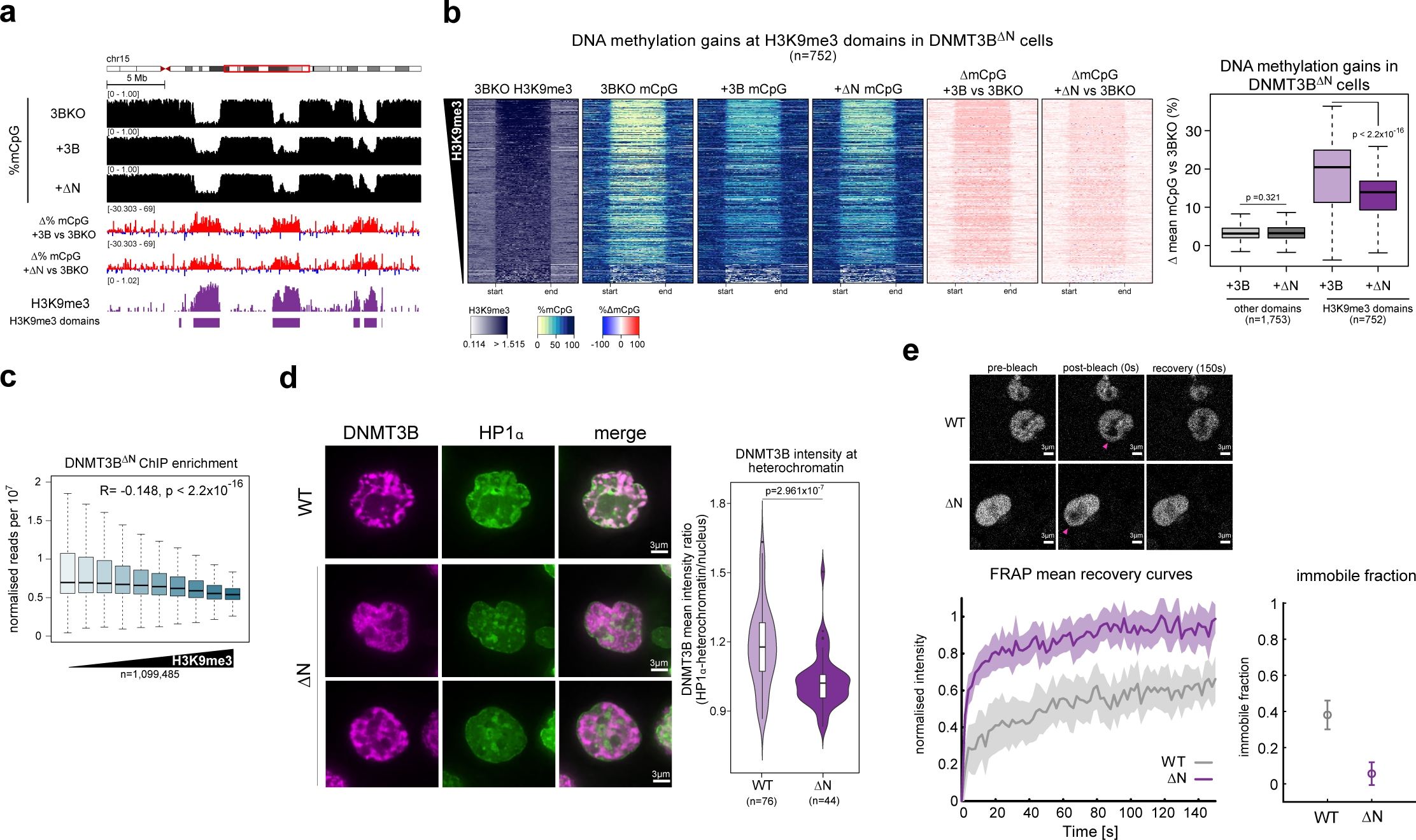
The N-terminus facilitates methylation of heterochromatin by DNMT3B. **a, b)** Lower DNA methylation recovery at heterochromatin upon expression of DNMT3B lacking the N-terminal region compared to full length DNMT3B. **a)** Representative genomic location showing gains of DNA methylation at H3K9me3 domains in DNMT3BKO cells expressing DNMT3B or DNMT3B^ΔN^. Genome browser plots show absolute (black) and differential (gain=red, loss=blue) DNA methylation levels, DNMT3B KO ChIP-seq signals and H3K9me3 domains defined in DNMT3B KO cells. ChIP-seq are normalised reads per 10^6^. **b)** Left, heatmaps showing levels of H3K9me3, of absolute and differential DNA methylation at DNMT3B KO H3K9me3 domains in DNMT3B^WT^ or DNMT3B^ΔN^ cells. The domains are ranked by their mean H3K9me3 levels. Right, boxplot showing gains of DNA methylation at H3K9me3 domains or domains not marked by neither H3K9me3 nor H3K27me3 DNMT3B^WT^ or DNMT3B^ΔN^ cells. All measurements for DNMT3B^ΔN^ are mean of two independently generated cell lines. **c, d)** Decreased DNMT3B localisation at heterochromatin in the absence of the NTD. **c)** Boxplot showing T7-DNMT3B^ΔN^ enrichment normalised over input at 2.5 kb genomic windows of increased H3K9me3 enrichment in DNMT3B KO cells grouped into deciles. Pearson’s correlation coefficient (R) is shown alongside its associated p-value. **d)** Left, representative confocal images of SNAP-DNMT3B and GFP-HP1⍺ localisation in live DNMT3B KO cells. Bars= 3µm. Right, violin plot showing the distribution of DNMT3B mean intensity ratio between HP1⍺-marked heterochromatin and the rest of the nucleus from one representative experiment. P-value is from two-sided Wilcoxon rank sum test. DNMT3B^WT^ data are repeated from Fig. 3e as the data were part of the same experiment. **e)** Top, representative images from a FRAP experiment (pink arrow indicates the side of the bleached area). Bottom left, mean fluorescence recovery curves for DNMT3B^WT^ and DNMT3B^ΔN^ from three independent FRAP experiments. Propagated error is shown by the shaded area. Bottom right, mean immobile fraction of DNMT3B^WT^ and DNMT3B^ΔN^ calculated from 100 to 150 s for three independent experiments. Bars represent the propagated error from the standard deviations. For all boxplots: lines = median; box = 25th–75th percentile; whiskers = 1.5 × interquartile range from box and p-values are from two-sided Wilcoxon rank sum tests.

We then assessed DNMT3B^ΔN^ localisation on chromatin using ChIP-seq. DNMT3B^ΔN^ signal was significantly correlated with H3K36me3 levels across the genome, similarly to DNMT3B^WT^ (*Fig. S6e* and *f*). However, DNMT3B^ΔN^ signal had a stronger negative correlation with H3K9me3 than DNMT3B^WT^ (Pearson’s correlations, R=-0.148 and -0.083 respectively, both p < 2.2×10^-16^, *Fig. 6c*), consistent with deletion of the N-terminus disproportionately affecting DNMT3B localisation to heterochromatin.

Using the biscistronic fluorescent protein stability assay, we observed that the ratio of GFP-DNMT3B^ΔN^ to dsRed was similar to that of GFP-DNMT3B^WT^ protein (*Fig. S6g*), suggesting that removal of the N-terminus did not affect the stability of the protein. However, different expression levels of DNMT3B^ΔN^ and DNMT3B^WT^ in cell populations could account for the lower DNA methylation levels at heterochromatin in DNMT3B^ΔN^ cells. To independently test the role of DNMT3B’s N-terminal region in recruiting it to heterochromatin, we analysed the localisation of SNAP-tagged DNMT3B^ΔN^ in live cells using confocal microscopy. The ratio of DNMT3B signal at HP1*α* -marked heterochromatin versus the rest of nucleus was significantly lower for DNMT3B^ΔN^ than for DNMT3B^WT^, (*Fig. 6d* and *Fig. S6h*), supporting the hypothesis that removal of the N-terminal region affects the recruitment of DNMT3B to heterochromatin.

To confirm the N-terminal region contributes to DNMT3B interaction with chromatin we performed fluorescence recovery after photobleaching (FRAP) in cells expressing SNAP-DNMT3B^WT^ or DNMT3B^ΔN^ (*Fig. 6e*, top). Over the time course, DNMT3B^ΔN^ showed higher recovery over the time course (*Fig. 6e*, bottom left) and a significantly smaller immobile fraction (*Fig. 6e*, bottom right) than DNMT3B^WT^. This is consistent with the proportion of DNMT3B^ΔN^ with slow dynamics in cells being lower than for DNMT3B^WT^. It suggests that less of DNMT3B^ΔN^ is bound to chromatin than DNMT3B^WT^ and supports a role for the N-terminus in stabilising DNMT3B’s association with chromatin.

Taken together, these data suggest that the N-terminus of DNMT3B facilitates the PWWP-independent recruitment of DNMT3B to constitutive H3K9me3-marked heterochromatin.

## Discussion

Here we have analysed *DNMT3B* mutations and knockout human cells to show that mutations in DNMT3B’s PWWP domain result in hypermethylation of H3K9me3-marked heterochromatin loci. Our results demonstrate that DNMT3B is recruited to and methylates heterochromatin in a PWWP-independent manner that is facilitated by its N-terminal region. We propose that the hypermethylation of heterochromatin observed with PWWP mutations results from the redistribution of DNMT3B away from H3K36me3 marked loci (*Fig. 7*).

**Figure 7.**
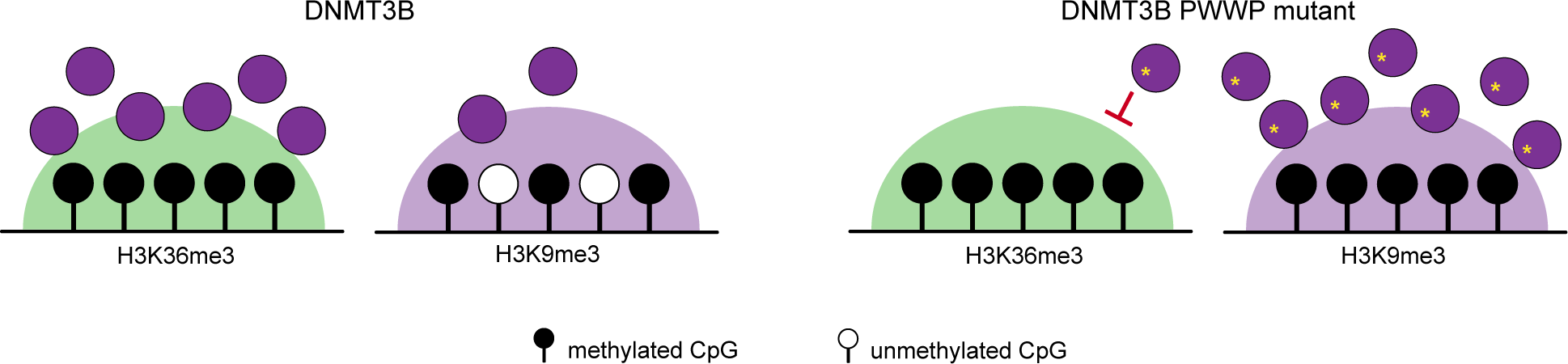
DNMT3B PWWP mutations cause hypermethylation of heterochromatin. Normally, DNMT3B predominantly localises to H3K36me3-marked regions through the action of its PWWP domain (Baubec et al., 2015). A small proportion of DNMT3B is also localised to H3K9me3-marked heterochromatin. When its PWWP is mutated, binding to H3K36me3-marked regions is inhibited and an increased pool of DNMT3B localises to heterochromatin in a manner that is facilitated by the N-terminal region resulting in gains of DNA methylation in this genomic compartment.

Previous work has shown that the majority of DNMT3B localises to actively transcribed gene bodies due to the interaction of the PWWP domain with H3K36me3 (Baubec et al., 2015) and that losses of methylation in *Dnmt3b* knockout mouse ES cells correlate with H3K36me3 (Neri et al., 2017). However, loss of DNMT3B has also been shown to result in hypomethylation of heterochromatic satellite loci in mouse (Okano et al., 1999) and human cells (Jeanpierre et al., 1993; Liao et al., 2015) and DNMT3B localises to heterochromatin blocks in interphase cells (Chen et al., 2004; Ge et al., 2004). In these studies, mutations of DNMT3B’s PWWP domain were reported to abrogate the localisation of DNMT3B to these heterochromatic domains, but the effect of the mutations on protein stability was not tested. We instead show that multiple, stable mutations affecting the interaction of the PWWP domain with H3K36me3 or DNA result in increased activity of DNMT3B at heterochromatin.

Our results parallel gains of methylation at DNA methylation valleys reported for PWWP mutations affecting the interaction of DNMT3A with methylated H3K36 in HESJAS patients (Heyn et al., 2019) and a mouse model (Sendžikaitė et al., 2019). Recent papers have proposed that this re-localisation is due to the recognition of the Polycomb Repressive Complex 1-associated histone mark H2AK119ub by DNMT3A’s N-terminus (Gu et al., 2022; Weinberg et al., 2021). The N-terminal region of DNMT3B remains less well understood but it has been reported to facilitate nucleosome binding in cells (Jeong et al., 2009). Future studies are likely to further illuminate the role of DNMT3B’s N-terminus in recruiting it to the genome. A previous report suggested that DNMT3B interacts with HP1⍺ and is recruited to pericentromeric repeats in mouse cells in a Suv39h dependent manner (Lehnertz et al., 2003). However, they did not define which regions of DNMT3B were responsible for localisation to heterochromatin and did not analyse the non-repetitive genome. A recent study suggests that the HESJAS causing mutation W330R (Heyn et al., 2019) may also mis-localise DNMT3A by promoting DNA binding (Lue et al., 222022). This is unlikely to explain our results with DNMT3B, as while we have analysed a mutation of a paralogous residue (W263A), we observe similar results with multiple mutations and deletion of the PWWP domain. While some of the mutations we studied do not directly affect DNMT3B’s interaction with H3K36me3, it has previously been suggested that high-affinity association of PWWP domains with chromatin requires cooperative DNA and histone-mark binding (Eidahl et al., 2013; van Nuland et al., 2013; Wang et al., 2020). It is also possible that other parts of DNMT3B play a role in its action at heterochromatin, for example a recent study suggests that the ADD domain of DNMT3B behaves differently to that of DNMT3A (Boyko et al., 2022).

Recessive mutations of DNMT3B cause ICF1 syndrome (Weemaes et al., 2013), a Mendelian condition characterised by hypomethylated heterochromatin (Heyn et al., 2012; Jeanpierre et al., 1993). The majority of mutations truncate DNMT3B or reduce its catalytic activity (Weemaes et al., 2013). A recent study also suggested that some ICF1 mutations can affect the formation of DNMT3B homo-oligomers (Gao et al., 2022). However, two patients have been described as carrying the S270P mutation which lies within the PWWP domain (Shirohzu et al., 2002). This mutation has been shown to reduce the binding of the domain to H3K36me3 (Baubec et al., 2015). Differences in methylation in gene bodies marked by H3K36me3 have also been proposed to lead to altered splicing in ICF1 patient cells (Gatto et al., 2017). Our results show that S270P destabilises DNMT3B *in vitro* and *in vivo* explaining why it does not cause hypermethylation of heterochromatin like the other PWWP mutations we have analysed. We also suggest that the pathogenic nature of S270P in causing ICF1 syndrome is because it reduces DNMT3B levels in cells rather than its effect on association with H3K36me3.

Loss of DNA methylation at non-repetitive heterochromatic loci is a hallmark of cancer and aging and results in the formation of partially methylated domains (Zhou et al., 2018). Our results here and those of our previous study of CpG islands (Masalmeh et al., 2021) demonstrate that the majority of DNMT3B is localised to H3K36me3 in cancer cells. However, alterations in the balance of DNMT3B recruitment to H3K36me3 and heterochromatin could potentially play a role in aging and cancer-associated hypomethylation.

In conclusion, we report of an unexpected consequence of DNMT3B PWWP mutations which reveal that DNMT3B plays a role in DNA methylation homeostasis at heterochromatin, a genomic compartment whose DNA methylation levels are frequently altered in human disease.

## Materials and Methods

### Cell culture

HCT116 and DNMT3B KO cells were gifts from B. Vogelstein (Rhee et al., 2002). Cells were cultured in McCoy’s 5A medium (Gibco) supplemented with 10% fetal calf serum (Life technologies) and penicillin-streptomycin antibiotics at 140 and 400 μg/ml respectively.

### Generation of CRISPR/Cas9-edited HCT116 cell lines

To introduce the W263A mutation in endogenous *DNMT3B* in HCT116 cell, the CRISPR Targets track from the USCS genome browser (https://genome.ucsc.edu) was used for guide RNA design and the corresponding oligonucleotides cloned into pSpCas9(BB)-2A-GFP (pX458, Addgene Plasmid 48138, a gift of F. Zhang). The donor template for homology-directed repair was an 80 nt ss-oligo Alt-R HDR Donor Oligo (Integrated DNA Technologies) designed on exon 7 and carrying the W263A mutation. HCT116 cells were transfected with the vector and ss-oligo, and GFP +ve cells selected by FACS 48 h after transfection and plated at clonal density. Individual colonies were screened by PCR followed by Sanger sequencing. Primers and oligonucleotides used are listed in *Supplementary Table 1*.

### Generation of plasmid constructs

To create piggyBac DNMT3 expression vectors, DNMT3 sequences from pcDNA3-Myc-DNMT3B2 and pcDNA3-Myc-DNMT3A1 plasmids (Addgene plasmids 36942 and 35521, a gift from A. Riggs) (Chen et al., 2005) were initially subcloned into the pCG plasmid (a gift from N. Gilbert) downstream of the T7 tag. The catalytically inactive point mutations C631S (DNMT3B2) and C710S (DNMT3A1) and the point mutations in the PWWP domain of DNMT3B (W263A, D266A, S270P, K276E and K294E) were introduced using the QuikChange II site-directed mutagenesis kit (Agilent). DNMT3B^ΔN^ starts from M200 and was generated by PCR. DNMT3B^ΔPWWP^ was synthesised using gBlocks (Integrated DNA Technologies) and generates a DNMT3B truncated from E206 to Y375. T7-tagged DNMT3s were then cloned into PB-CGIP (a gift from M. McCrew) (Macdonald et al., 2012) by swapping eGFP. To create vectors for live-cell imaging a SNAP-tag was initially subcloned from pCS2-SNAP (a gift from D. Papadopolous) to pB530A-puroVal2, previously created by substituting the copepod GFP from the pB530A plasmid (System Biosciences) with the Puromycin resistance gene. DNMT3B and mutants sequences were cloned downstream the SNAP-tag. To generate vectors for the protein stability assay DNMT3B and mutant sequences were cloned into pLenti-DsRed-IRES-EGFP (Addgene plasmid 92194, a gift from Huda Zoghbi).

### DNMT3 expression in cells

HCT116 and DNMT3B KO cells were transfected with FuGENE HD transfection reagent (Promega). For stable integrants, DNMT3 expression constructs were co-transfected with a plasmid expressing piggyBac transposase. After 48 h cells stably expressing DNMT3s were selected with 1 μg/ml puromycin and expanded in the presence of puromycin for 3-4 weeks before being harvested for analysis.

### DNA extraction

To extract genomic DNA cells were resuspended in genomic lysis buffer (300 mM NaCl, 1% SDS, 20 mM EDTA) and incubated with Proteinase K (Roche) at 55 °C overnight. RNA was removed by incubation with RNase A/T1 Cocktail (Ambion) at 37°C for 1 h, in between two phenol-chloroform extraction steps. DNA was quantified by Nanodrop 8000 spectrophotometer and purity assessed by electrophoresis.

### Global measurement of DNA methylation by mass-spectrometry

DNA for mass-spectrometry and following analysis was performed as previously described (Masalmeh et al., 2021). 1 μg genomic DNA was denatured at 95 °C for 10 min in 17.5 μl water. DNA was then digested to nucleotides overnight at 37 °C with T7 DNA polymerase (Thermo Scientific). The reaction was inactivated by incubating at 75 °C for 10 min. Samples were then centrifuged for 45 min at >12,000 g and the supernatant transferred into new tubes for analysis. Enzyme was removed by solvent precipitation. The samples were adjusted back to initial aqueous condition and volume and LC-MS was performed on a Thermo Ultimate 3000/ Thermo Q Exactive system, using a Hypercarb 3 μm × 1 mm × 30 mm Column (Thermo 35003-031030) and gradient from 20 mM ammonium carbonate to 2 mM ammonium carbonate 90% acetonitrile in 5 min. Data were acquired in negative mode, scanning at 70 k resolution from 300 to 350 m/z. Extracted ion chromatograms were analysed using Xcalibur (Thermo Scientific, v2.5-204201/2.5.0.2042) to extract peak intensities at the m/z values expected for each nucleotide (based on annotation from METLIN) (Smith et al., 2005) following manual inspection to ensure that they were resolved as clear single peaks. The percentage of 5-methylcytosine present in the sample was calculated as the ratio of the area under the 5-methylcytosine peak to the area under the guanine peak.

### WGBS data generation

100 or 200 ng of purified DNA samples were sheared using the Covaris E220 Evolution Focused Ultrasonicator to create fragments of approximately 350 bp. 0.5 ng of unmethylated phage-λ DNA (NEB) was spiked into each DNA sample prior to shearing to allow assessment of the efficiency of the bisulfite-conversion reaction. 100 ng of each sheared DNA sample was then processed using the EZ DNA Methylation Gold Kit or EZ DNA Methylation-Lightning Kit (Zymo Research) according to the manufacturer’s protocol to create bisulfite-converted single-stranded (BC-ss) DNA. For WGBS experiments in *Fig. 1* and *S1* comparing DNMT3B KO cells and reintroduction of DNMT3B, the TruSeq DNA Methylation Kit (Illumina Inc.) was used. Bisfulfite converted single-stranded DNA was randomly primed with a polymerase able to read uracil nucleotides to synthesise DNA strands containing a sequence specific tag. 3’ ends of the newly synthesized DNA strands were then selectively tagged with a second specific sequence tag to produce di-tagged DNA molecules with known sequence tags at 5’ and 3’ ends. Di-tagged molecules were then enriched by PCR with unique indexed PCR primers to provide dsDNA libraries with Illumina sequencing adapters that could be multiplexed and sequenced on a single flow cell. For all other WGBS experiments the Accel-NGS Methyl-Seq DNA Library Kit (Swift BioSciences) was used. For the Swift kit, bisfulfite converted single-stranded DNA was first denatured before simultaneous tailing and ligation of truncated adapters to 3’ ends with Adaptase. An extension step then incorporates the truncated adapter 1 by a primer extension reaction, before a ligation step adds truncated adapter 2 to the bottom strand only. 9 cycles of PCR were used to increase yield and incorporate full-length adapters for unique dual-indexed sequencing. Finally, Agencourt AMPure XP beads (Beckman Coulter) were used to remove oligonucleotides and small fragments and change enzymatic buffer composition. Libraries were quantified using the Qubit dsDNA HS assay kit, assessed for size and quality using the Agilent Bioanalyser with the DNA HS Kit and combined in a single equimolar pool. Sequencing was performed on a P2 flow cell on the Illumina NextSeq 2000 platform using NextSeq 1000/2000 P2 Reagents v3 kit (200 Cycles), or on the NextSeq 550 platform using NextSeq 500/550 High-Output v2 kit (150 cycles). PhiX Control v3 Library was spiked in at a concentration of ∼5% to increase library diversity for sequencing. Library preparation and sequencing was performed by the Edinburgh Clinical Research Facility.

### WGBS data processing

Sequencing quality was assessed with FASTQC (*v0.11.4*). Low quality reads and remaining adaptors were removed using TrimGalore (*v0.4.1*, Settings: *--adapter AGATCGGAAGAGC --adapter2 AAATCAAAAAAAC*). The paired end reads were then aligned to the hg38 genome using Bismark (*v 0.18.1* with Bowtie2 *v2.3.1* and settings: *-N 0 -L 20*, trimming settings *--clip_r1 10 --clip_r2 13 -- three_prime_clip_r1 1 --three_prime_clip_r2 1*) (Krueger and Andrews, 2011; Langmead and Salzberg, 2012) before PCR duplicates were identified and removed using Bismark’s *deduplicate_bismark* command. Aligned BAM files were processed to report coverage and number of methylated reads for each CpG observed. Forward and reverse strands were combined using Bismark’s *methylation extractor* and *bismark2bedgraph* modules with custom Python and AWK scripts. Processed WGBS files were assessed for conversion efficiency based on the proportion of methylated reads mapping to the phage-λ genome spike-in (>99.5% in all cases). For summary of WGBS alignment statistics see *Supplementary Table 2*. BigWigs for visualisation of WGBS data were generated using the coverage in 2.5 kb-sized windows across the genome. These were defined using BEDtools (*v2.27.1*) (Quinlan and Hall, 2010) and coverage and converted to bigWigs using UCSC tools *bedGraphToBigWig* (*v326*). Windows with a total coverage <5 were excluded before conversion to bigWigs.

### WGBS data analysis

WGBS data were analysed by defining genomic windows for analysis. For genome-wide analyses, non-overlapping 2.5 kb-sized genomic windows were defined using BEDtools *makewindows* (*v2.27.1*). The percentage methylation within these windows was calculated as the weighted mean methylation using the observed coverage of CpGs within each window (unconverted coverage/total coverage) using BEDtools. Windows with a total coverage <10 were excluded from the analysis.

To analyse methylation within domains, BEDtools was used to calculate the weighted mean coverage from CpGs located within the domain as defined above. To analyse the profile around domains, heatmaps were generated by defining 40 scaled windows across each domain and 20 windows of 25 kb upstream and downstream of each domain relative to the forward strand using custom R scripts. The weighted mean coverage in each window was then defined as above using BEDtools. Domains were excluded from analysis if ≥ 80% of the 40 scaled windows across the domain had a coverage of 0 in all samples in the analysis.

For the methylation profile across genes, a similar analysis was performed. Transcript locations were downloaded from ENSEMBL (*v106*). Only coding transcripts were considered (those defined as ‘protein_coding’ by ENSEMBL) and each gene’s weighted mean methylation was defined from the mean of all coding transcripts annotated to it. To analyse the profile around genes, heatmaps were generated by defining 40 scaled windows across each coding transcript and 20 windows of 250 bp upstream and downstream of each coding transcript relative to its direction of transcription using custom R scripts. Before calculating the mean profile for each gene, transcripts were excluded if ≥ 80% of the 40 scaled windows across the domain had a coverage of 0 in all samples in the analysis.

To statistically test differences in DNA methylation levels, differential mean methylation percentage across domains were compared using a Wilcoxon rank sum test.

### ChIP-seq data generation

ChIP-seq was performed as previously described (Masalmeh et al., 2021). For T7-DNMT3B ChIP-Rx-seq experiments, 1×10^7^ cells were harvested, washed and crosslinked with 1% methanol-free formaldehyde in PBS for 8 min at room temperature. Crosslinked cells were lysed for 10 min on ice in 50 μl of lysis buffer (50 mM Tris-HCl pH 8, 150 mM NaCl, 1 mM EDTA, 1% SDS) freshly supplemented with proteinase inhibitor (Sigma-Aldrich). IP dilution buffer (20 mM Tris-HCl pH 8, 150 mM NaCl, 1 mM EDTA, 0.1% Triton X-100) freshly supplemented with proteinase inhibitor, DTT and PMSF was added to the samples to reach a final volume of 500 μl. As prolonged sonication caused T7-DNMT3B degradation, chromatin was fragmented using Benzonase (Pchelintsev et al., 2016): samples were sonicated on ice with Soniprep 150 twice for 30 s to break up nuclei; then 200 U of Benzonase Nuclease (Sigma) and MgCl_2_ (final concentration 2.5 mM) were added and samples were incubated on ice for 15 min. The reaction was blocked by adding 10 μl of 0.5 M EDTA pH 8. Following centrifugation for 30 min at 18,407 g at 4 °C, supernatants were collected and supplemented with Triton X-100 (final concentration 1%) and 5% input aliquots were retained for later use. Protein A Dynabeads (Invitrogen) previously coupled with 10 μl of T7-Tag antibody per 1×10^7^ cells in blocking solution (1x PBS, 0.5% BSA) were added and the samples incubated overnight under rotation at 4 °C. Beads were then washed for 10 min at 4 °C with the following buffers: IP dilution buffer 1% Triton X-100 (20 mM Tris-HCl pH 8, 150 mM NaCl, 2 mM EDTA, 1% Triton X-100), buffer A (50mM HEPES pH 7.9, 500 mM NaCl, 1 mM EDTA, 1% Triton X-100, 0.1% Na-deoxycholate, 0.1% SDS), buffer B (20 mM Tris pH 8, 1 mM EDTA, 250 mM LiCl, 0.5% NP-40, 0.5% Na-deoxycholate), TE buffer (1mM EDTA pH 8, 10 mM Tris pH 8). Chromatin was eluted by incubating the beads in extraction buffer (0.1 M NaHCO_3_, 1% SDS) for 15 min at 37 °C. To reverse the cross-linking Tris-HCl pH 6.8 and NaCl were added to final concentrations of 130 mM and 300 mM respectively, and immunoprecipitations were incubated at 65 °C overnight.

Samples were then incubated at 37 °C for 1 h after addition of 2 μl of RNase Cocktail Enzyme Mix (Ambion). Then 40 μg of Proteinase K (Roche) were added, followed by 2 h incubation at 55 °C. Input material was similarly de-crosslinked. Samples were purified with the MinElute PCR purification kit (QIAGEN). For ChIP-Rx-seq of endogenous T7-DNMT3B and ectopic T7-DNMT3B expressed in DNMT3B KO cells, 20 μg of Spike-in chromatin (ActiveMotif 53083) was added to each sample after sonication. 2 μl of spike-in antibody per sample (ActiveMotif 61686) was also added in a ratio 1:5 versus the T7 antibody. A similar protocol was used for H3K4me3, H3K9me3, H3K27me3 and H3K36me3 ChIP-seq experiments, except: 0.5×10^7^ cells were harvested and crosslinked with 1% methanol-free formaldehyde in PBS for 5 min at room temperature. For H3K36me3 ChIP-Rx-seq crosslinked Drosophila S2 cells were spiked into samples before sonication at a ratio of 20:1 human to Drosophila cells. Following nuclei rupture by sonication on ice with Soniprep 150, chromatin was fragmented using Bioruptor Plus sonicator (Diagenode) for 40 cycles (30 s on/30 s off on high setting at 4 °C). 2 μl /1×10^6^ cells of the following antibodies were used for immunoprecipitations: H3K4me3 (EpiCypher 13-00041), H3K9me3 (Active Motif 39161), H3K27me3 (Cell Signaling Technology C36B11) and H3K36me3 (Abcam ab9050). Libraries were prepared using the NEBNext Ultra II DNA Library Prep Kit for Illumina (E7645) according to the manufacturer instructions. NEBNext Multiplex Oligos for Illumina (NEB) barcode adapters were used. Specifically, Illumina Index Primers Set 1 (E7335) for endogenous T7-DNMT3B and H3K36me3, and Unique Dual Index UMI Adaptors DNA Set 1 (E7395) for the other libraries. For histone modifications ChIP-seq, adapter-ligated DNA was size selected for an insert size of 150 bp using Agencourt AMPure XP beads. Libraries were quantified using the Qubit dsDNA HS or BR assay kit and assessed for size and quality using the Agilent Bioanalyser. Endogenous T7-DNMT3B and H3K36me3 ChIP-Rx-seq libraries were sequenced using the NextSeq 500/550 high-output version 2.5 kit (75 bp paired end reads, DNMT3B or 75 bp single end reads, H3K36me3). The other ChIP-seq libraries were sequenced using the NextSeq 2000 P3 (50 bp paired end reads). Libraries were combined into equimolar pools to run within individual flow cells. Sequencing was performed by the Edinburgh Clinical Research Facility.

### ChIP-seq data processing

All ChIP-seq experiments were processed previously described (Masalmeh et al., 2021). Read quality was checked using FASTQC (*v0.11.4*, https://www.bioinformatics.babraham.ac.uk/projects/fastqc), with low quality reads and adaptors removed using TrimGalore with default settings (*v0.4.1*). Reads were aligned to hg38 using bowtie 2 (*v2.3.1*, with settings: *-N 1 -L 20 --no-unal*) (Langmead and Salzberg, 2012). For paired end data, additional settings were used during alignment to remove discordant reads: *--no-mixed –nodiscordant -X 1000*. ChIP-Rx-seq reads were aligned to a combination of hg38 and dm6 genomes. Multimapping reads excluded using SAMtools (*v1.6*, with settings: *-bq 10*) (Li et al., 2009) and PCR duplicates excluded using SAMBAMBA (*v0.5.9*) (Tarasov et al., 2015). For T7 ChIP-Rx-seq UMIs were used for de-duplication instead of SAMBAMBA. We first extracted UMIs from the appropriate read and placed them in the read FASTQ headers using UMI tools *extract* (*v1.0.0* and setting: *--bc-pattern=NNNNNNNNNNN*). Then deduplication was performed using the UMI tools *dedup* function (*v1.0.0* and setting: *--paired*). For summary of ChIP-seq alignment statistics see *Supplementary Table 3*.

Tracks for visualisation of histone ChIP-seq were generated using Deeptools (*v3.2.0*) (Ramirez et al., 2016). Counts per million normalised tracks were generated using the *bamCoverage* function (settings: *--normalizeUsing CPM*) with the default bin size of 50 bp. The mean of replicate tracks and normalisation over the input was calculated using the *bigwigCompare* function (settings: *--operation mean*). For the single-ended H3K36me3 ChIP-seq, the estimated fragment length of 150 bp was used. For the paired-end knock-in T7-DNMT3B ChIP-Rx-seq from HCT116 cells, the actual fragment size was used. Tracks for visualisation of T7 ChIP-Rx-seq were generated as follows: fragment counts in 2.5 kb sized windows were generated using BEDtools, normalised over input and scaled according to the spike-in (see ChIP-seq data analysis for details). The bedgraphs were then converted into bigwigs using UCSC tools *bedGraphToBigWig* (*v326*).

### ChIP-seq data analysis

For ChIP-seq data analyses, data were analysed by defining genomic windows for analysis as previously described (Masalmeh et al., 2021). Non-overlapping 2.5 kb-sized genomic windows were defined using BEDtools *makewindows* (*v2.27.1*). Histone modifications and T7-DNMT3B normalised coverage in these genomic windows or domains were derived from ChIP-seq by first counting the number of reads or fragments overlapping the windows/domains using BEDtools *coverage* function. For paired end data the BAM file was first converted to a BED file of fragment locations using BEDtools *bamtobed* function. Coverage counts were scaled to counts per 10 million based on total number of mapped reads per sample and divided by the input read count to obtain a normalised read count. An offset of 0.5 was added to all windows prior to scaling and input normalisation to prevent windows with zero reads in the input sample generating a normalised count of infinity. Regions where coverage was 0 in all samples were removed from the analysis. We also excluded windows overlapping poorly mapped regions of the human genome (https://github.com/Boyle-Lab/Blacklist) (Amemiya et al., 2019) using BEDtools *intersect*. Gaps and centromeres identified using annotations from the UCSC browser (hg38 gap and centromere tracks) were similarly removed. First, annotations were downloaded from the UCSC table browser and regions annotated as heterochromatin, short arm and telomeres from the gaps track were merged with the centromeres track using BEDtools *merge* with - *d* set to 10 Mb. Windows overlapping this merged file were then excluded from the analysis using BEDtools *intersect*.

For ChIP-Rx-seq, before proceeding with the analysis, normalised coverage values were scaled using a scaling factor generated from the number of reads mapping to the *D.melanogaster* genome as previously described (Masalmeh et al., 2021). Reads mapping to the *D.melanogaster* genome in each ChIP and input sample were first scaled to reads per 1×10^7^. The scaling factor was then calculated as the ratio of the scaled *D.melanogaster* reads in two ChIP samples over their respective ratio from the input samples (modified from published method to take account of the presence of an input sample (Orlando et al., 2014)). This analysis was applied to each biological replicate and where multiple replicates were available, the mean was calculated for each window.

The profile around defined heterochromatic domains was similarly calculated but using 40 scaled windows across each domain and 20 windows of 25 kb upstream and downstream of each domain relative to the forward strand using custom R scripts. Scaled normalised coverage values were calculated as above for genomic windows except that domains were excluded from analysis if ≥ 80% of the 40 scaled windows across the domain had a coverage of 0 in all samples in the analysis. The mean profile was calculated from these values using R. For the ChIP-seq profile across genes, a similar analysis was performed. Transcript locations were downloaded from ENSEMBL (*v106*). Only coding transcripts were considered (those defined as ‘protein_coding’ by ENSEMBL) and each gene’s scaled normalised coverage calculated from the mean of all coding transcripts annotated to it. To analyse the profile around genes, heatmaps were generated by defining 40 scaled windows across each coding transcript and 20 windows of 250 bp upstream and downstream of each coding transcript relative to its direction of transcription using custom R scripts. Scaled normalised coverage values were calculated as above for genomic windows except that transcripts were excluded from analysis if ≥ 80% of the 40 scaled windows across the domain had a coverage of 0 in all samples in the analysis. Colour scales for ChIP-seq heatmaps range from the minimum to the 90% quantile of the normalised read count.

### Definition and comparison of heterochromatic domains

H3K9me3 and H3K27me3 domains were called using a hidden Markov model as previously described (Spracklin and Pradhan, 2020). Normalised mean H3K9me3 and H3K27me3 ChIP-seq coverage in 25 kb genomic windows was analysed using the *bigwig_hmm.py* script (https://github.com/gspracklin/hmm_bigwigs) to define two states *(-n 2*). We excluded poorly mapped regions of the genome from these domains using annotations of gaps and centromeres from the UCSC browser (hg38 gap and centromere tracks). Annotations were downloaded from the UCSC table browser. Regions annotated as heterochromatin, short arm and telomeres from the gaps track were merged with the centromeres track using BEDtools *merge* with *-d* set to 10 Mb. This merged file was then excluded from the domains BED files using BEDtools *subtract*. H3K27me3 only domains were then defined as those which did not overlap a H3K9me3 domain using BEDtools intersect.

Comparisons between heterochromatic domains were conducted using BEDtools *jaccard* and *fishers* modules. PMDs were defined in HCT116 cells using methpipe (*v5.0.0*) (Decato et al., 2020). WGBS BAM files generated by Bismark before being processed and de-duplicated using methpipe. PMDs were then called using the *pmd* module from methpipe and the recommended settings (bin size was determined as 8 kb by methpipe using the setting *-i 1000*). We excluded poorly mapped regions of the genome from PMDs as for the H3K9me3 and H3K27me3 domains. We also excluded PMDs shorter than 200 kb. We downloaded a BED file of regions resistant to nuclease digestion in HCT116 cells from the 2-state definitions in Gene Expression Omnibus accession GSE135580 (Spracklin and Pradhan, 2020).

### Bisulfite PCR data generation

500 ng of genomic DNA was bisulfite converted with the EZ DNA Methylation-Gold kit (Zymo Research). Bisulfite PCR was performed using EpiTaq (Takara) using custom made locus specific primers containing 4 bp unique molecular identifiers (UMI) on each side of the amplicon and partial adapter sequences to enable library amplification. Primer sequences are listed in *Supplementary Table 1*. Libraries for sequencing were amplified using unique dual index Illumina adapters and NEBNext Ultra II Q5 Master Mix (NEB). One purification step with Agencourt Ampure XP beads was performed before and after library amplification. Sequencing was performed on the Illumina iSeq 100 System using the iSeq 100 i1 Reagent v2 (300 cycle) Kit. PhiX Control v3 (Illumina) was spiked into the run at a concentration of 30% to improve cluster resolution and enable troubleshooting in the event of a run failure.

### Bisulfite PCR data analysis

To analyse bisulfite PCR data we calculated the mean percentage methylation for each sequenced fragment from its 2 reads. Following demultiplexing, we removed the Illumina indexes from the FASTQ headers to ensure compatibility with UMI tools (Smith et al., 2017). The extract function of UMI tools (*v1.0.0* with setting *--extract-method=string --bc-pattern=NNNN --bc-pattern2=NNNN*) was then used to remove the UMIs from these FASTQ sequences and place them in the header. The 1-colour chemistry used in the iSeq results in the generation of artefactual high-confidence poly-G reads (https://sequencing.qcfail.com/articles/illumina-2-colour-chemistry-can-overcall-high-confidence-g-bases). A custom command line script was therefore used to identify and filter out paired reads where one, or both, of the reads had a guanine content of at least 90%. The remaining reads were trimmed using the *paired* setting of TrimGalore (*v0.6.6* with the setting: *--paired*) and aligned to the reference genome using Bismark (*v0.22.3* with settings: *--multicore 3 -N 0 -L 20*), indexed using Samtools (*v1.13*) and deduplicated using the *dedup* functionality of UMI tools (*v1.0.0* with settings: *--paired -- method=unique*). We then calculated the mean methylation of each fragment using a custom command line script. Briefly, Samtools was used to sort deduplicated reads by read name and the *bismark_methylation_extractor* function of Bismark (*v0.22.3* with settings: *-p --no_header -- no_overlap*) was used to extract the methylation state of CpGs on individual reads. The command line was then used to identify the number of called CpGs per read (n_r_) and the number of CpGs with a methylated call per read (m_r_), and this information was used to calculate an overall mean methylation level (m_r_/n_r_) for each read. Read statistics are provided in *Supplementary Table 4*.

### Methylation sensitive Southern blot

1.5 μg of purified genomic DNA was digested with the methylation sensitive BstBI enzyme (NEB) overnight and separated on 0.8% agarose electrophoresis gel. The gel was incubated in the following solutions: denaturation (0.5 M NaOH, 1.5 M NaCl), neutralisation (0.5 M Tris pH 7.5, 1.5 M NaCl) and 20x SSC, before the DNA was transferred to Hybond N+ membrane (Amersham). DNA was crosslinked to the membrane by UV irradiation (254 nm at 0.15 J), followed by hybridization in DIG Easy Hyb buffer (Roche) with satellite II DIG-labelled probe at 42 °C for at least 4 h. The probe was generated by PCR amplification of satellite II sequence from plasmid p375M2.4 (Jackson et al., 1992) (a gift from N. Gilbert) using M13 universal primers. The blot was then washed in low (2x SSC, 0.1% SDS) and high (0.5x SSC, 0.1% SDS) stringency buffers, prior to incubation with anti-digoxigenin-AP antibody (Roche). Detection was performed using CSPD chemiluminescent substrate (Roche) on a ImageQuant LAS4000. Quantification was performed using Fiji measuring the intensity of the top 1/10^th^ of each lane (undigested methylated DNA) normalised over the intensity of the bottom band indicated in the figures (digested hypomethylated DNA).

### Western blotting

Whole-cell extracts were obtained by sonication in urea buffer (8 M Urea, 50 mM Tris pH 7.5, 150 mM β-mercaptoethanol) and quantified using Pierce BCA Protein Assay Kit (Thermo Scientific). Extracts were analysed by SDS–polyacrylamide gel electrophoresis using 4-12% Bis-Tris NuPAGE protein gels (Life Technologies) and transferred onto nitrocellulose membrane in 2.5 mM Tris-base, 19.2 mM glycine and 20% methanol. The primary antibodies used are: T7-Tag (D9E1X, Cell Signalling Technology), DNMT3B (D7O7O, Cell Signalling Technology), GAPDH (14C10, Cell Signalling Technology), alpha-Tubulin (T6199, Sigma). Images were acquired with ImageQuant LAS4000 or LAS800 following incubation with HRP-conjugated IgG (Invitrogen). For western blots of CRISPR clones, IRDye secondary antibodies (LI-COR) were used and images acquired on The Odyssey CLx (LI-COR) machine.

### In cell fluorescent protein stability assay

DNMT3B KO cells were transfected with a polycistronic vector expressing dsRed and GFP-DNMT3Bs, generated by modifying pLenti-DsRed-IRES-EGFP (Addgene plasmid 92194, a gift from Huda Zoghbi), or control plasmids expressing either GFP or dsRed, or no plasmids. After 48 h cells were analysed by flow cytometry on BD LSR Fortessa Cell Analyser. Matrix compensation was set-up using negative (no plasmid), dsRed only and GFP only cells. FlowJo (BD Biosciences) was used to export the fluorescence intensities of individual dsRed+ cells. We then calculated the ratio for each cell and the mean ratio for each experiment using R. Each mutant was assessed in 3 independent transfection experiments.

### Protein purification

DNMT3B1 sequences equating to residues 213-351 (PWWP alone) or 213-555 (PWWP-ADD) were cloned into pLIC-6xHis-MBP-TEV vectors using ligation independent cloning. Sites were mutated either using site directed mutagenesis or direct cloning of synthesised double-strand gBlock fragments containing mutations (Integrated DNA technologies). DNMT3B constructs were expressed by induction at OD of 0.6-0.8 with 400 μM IPTG in BL-21 DE3 RIL *E.coli* overnight at 18 °C in 2x YT broth. Cell pellets were resuspended in lysis buffer (30 mM Sodium Phosphate pH 7.6, 400 mM NaCl, 10% glycerol, 0.1% Triton X-100, 1 μM ZnCl_2_, 1x Protease Inhibitor [284 ng/ml leupeptin, 1.37 μg/ml pepstatin A, 170 μg/ml phenylmethylsulfonyl fluoride and 330 μg/ml benzamindine], 1 mM AEBSF, 2 mM β-mercaptoethanol). For lysis the solution was supplemented with lysozyme (500 μg/ml), 2 mM MgCl_2_, and DNase (5 μg/ml) and sonicated. Clarified lysate was applied to pre-equilibrated bed of Ni-NTA Agarose beads (Qiagen), washed extensively with wash buffer (15 mM Sodium Phosphate pH 7.6, 500 mM NaCl, 10% glycerol, 2 mM β-mercaptoethanol, 30 mM Imidazole) followed by elution buffer (20 mM Tris pH 7.5, 400 mM NaCl, 300 mM Imidazole, 10% glycerol, 2mM β-mercaptoethanol). Protein containing fractions were further purified by size exclusion chromatography using Superdex 200 10/300 (GE Healthcare) gel-filtration column in SEC buffer (20 mM HEPEs pH 7.5, 150 mM NaCl, 5% glycerol, 2 mM DTT) and the main mono-disperse protein containing peak was collected, concentrated, flash frozen in liquid nitrogen and stored at -80 °C. Protein concentrations were determined via absorbance on a NanoDrop One spectrophotometer and subsequent SDS-PAGE with comparison to known amounts of control proteins (*Fig. S4c* and *S5a*) and protein stability of mutants was compared by thermal denaturation assay (see below).

### Thermal denaturation assay

Previously purified His-MBP-TEV-DNMT3B^213-555^ proteins were diluted to 1.5 μM in normalised TDA buffer (20 mM HEPES, 150 mM NaCl, 1 mM DTT, 0.5% glycerol, 1 μM ZnCl_2_, 5x SYPRO orange from Life Technologies). 50 μl reactions were imaged in triplicate in a 96-well plate. Thermal denaturation assays were performed in a Biometra TOptical RT-PCR machine, increasing temperature from 25 °C to 69.5 °C with 0.5 °C increments. Three measurements were taken for every step using an excitation wavelength of 490 nm and detection at 580 nm. Data was processed in instrument software and melting point derived the first derivative of the signal curve. Results from two biological replicates, each with measurements from three technical replicates were used to calculate mean T_m_ values and standard deviations.

### Multiple sequence alignment

The multiple sequence alignments shown in *Fig. 2a*, *5a* and *S4a* were performed with Clustal Omega using the following Uniprot sequences: Q9UBC3 (hDNM3B), O88509 (mDnmt3b), Q9Y6K1 (hDNMT3A) and O88508 (mDnmt3a). Colours represent the percentage of identity and were obtained using Jalview (*v2.11.2*) (Waterhouse et al., 2009).

### *In silico* analysis of PWWP domain and DNA binding residues

Building on previous studies reporting DNMT3B binding to DNA (Zhou et al., 2018), the structures of DNMT3B’s PWWP domains (mouse Protein Data Bank code 1KHC, (Zhou et al., 2018), human 5CIU (Rondelet et al., 2016)) were compared to homologous PWWP structures found to bind to DNA either in isolation (Protein Data Bank code 6IIT, (Tian et al., 2019), 7LH9 (Zhang et al., 2021)) or in a modified nucleosome core particle (LEDGF PDB 6S01, (Wang et al., 2020)). Rigid body modelling revealed conserved lysine and arginine residues as potential DNA binding residues. We identified two separable patches distal to the main H3 binding pocket, patch 1 comprising Lys-251 Arg-252 and patch 2 comprising Lys-276 Lys-232. We also observed a further two positively charged residues, Lys-234 and Lys-268. However, these were deemed to be too close to the methyl-lysine binding pocket of the PWWP to mutate without affecting methylated lysine 36 binding. Comparison and PWWP-domain figures were prepared in UCSF Chimera (Pettersen et al., 2004).

### Electrophoretic mobility shift assay for DNA binding

175 bp Widom-601 DNA fragments were generated by PCR based amplification as previously described (Wilson et al., 2019). A primer pair containing 5’ 6-FAM fluorophore on one oligo were synthesised and HPLC purified by Integrated DNA technologies (sequences atggaaCacatTGCACAGGATGTAT & /56-FAM/aATAgccacCtGCCCTGGAG). Large scale PCR of 384 100 μl reactions were pooled, filtered and purified using a ResourceQ column and eluted over a salt gradient. Fractions with appropriate DNA and fluorophore were pooled and ethanol precipitated prior to resuspension in TLE buffer (10 mM Tris pH 8, 0.1 mM EDTA). 3 nM of fluorescent 175 bp Widom-601 DNA was incubated with serial dilutions (from 4 μM to 62.5 nM) of recombinant 6xHis-MBP-DNMT3B^213-351^ and variants and in 15 μl of EMSA buffer (15 mM HEPES pH 7.5, 75 mM NaCl, 0.05 mg/ml BSA, 10% glycerol, 0.05% Triton X-100, 1 mM DTT, 8% sucrose, 0.01% bromophenol blue). Samples were incubated at 4 °C for 1 h to ensure end point of binding was reached. Products were separated on a native 5% polyacrylamide gel with 1x Tris-glycine running buffer for 2 h at 4 °C. Gels were imaged on ChemiDoc MP (BioRAD) using Alexa 488 exposure setting (blue epi illumination excitation and 532 nm/28 nm filter) to visualise fluorescent DNA. Gels were stained using Diamond DNA stain (Promega) and colloidal blue protein stain to confirm loading. EMSA assays were repeated, at least in triplicate.

### DNMT3B localisation data acquisition and image analysis

For live cell imaging DNMT3B KO cells were seeded in poly-L-lysine treated 8 well glass-bottom plates (ibidi GmBH) and transfected with SNAP-DNMT3B plasmid in addition to HP1⍺-GFP plasmid (a gift from Tom Misteli, Addgene plasmid 17652). After 48 h cells were incubated for 30 min at 37 °C with SNAP-Cell 647-SiR substrate (NEB, final concentration 1.2 µM). Cells were then incubated for 30 min at 37 °C in fresh media to wash off unbound substrate. Imaging was performed using FluoroBrite DMEM (Gibco) supplemented with 10% fetal calf serum. Imaging was carried out with a SoRa (Nikon-Europe) using a 100x 1.49NA oil immersion lens. Z-stack images were processed according to the manufacturers recommended settings using NIS-Elements AR software (*v5.1*, Nikon Europe).

To analyse the degree of DNMT3B localisation to heterochromatin, we used Imaris 9.8.0 (Bitplane, Oxford Instruments) to define heterochromatic regions of the nucleus using the ‘Surfaces’ function of Imaris to create a 3D isosurface of HP1*α*-GFP. The ‘Image Calculator’ Imaris XTension was used to create a channel that was the sum of both HP1*α*-GFP and the SNAP-tagged DNMT3B and the resultant channel was used to create a 3D isosurface representing the whole nucleus. This was done using the additive channel to ensure the whole nucleus was segmented. The Imaris module ‘Cell’ was then used to relate the segmented heterochromatin to the nucleus it was inside. This meant a ratio of the mean intensity of DNMT3B signal in the heterochromatic regions over the mean intensity in the rest of the nucleus could be obtained.

To analyse the degree of DNMT3B signal at the nuclear periphery from the same cells, we applied the ‘Distance Transformation’ XTension from Imaris to the nucleus isosurface to create a distance map channel where each voxel in the surface is valued by the distance in microns from the outside of the surface. By thresholding this channel from 0.1 – 0.5 a 3D isosurface representing the nuclear periphery was obtained. The mean intensity of this periphery has been represented as a ratio with the mean intensity of the rest of the nucleus. Z-planes used in figures were assembled and annotated using Fiji (https://fiji.sc).

### FRAP data acquisition and analysis

For FRAP, DNMT3B KO cells were seeded in poly-L-lysine treated 8 well glass-bottom plates (ibidi GmBH) and transfected with SNAP-DNMT3B plasmid. After 48 h cells were incubated for 30 min at 37 °C with SNAP-Oregon green (NEB, final concentration 4 µM). Cells were then incubated for 30 min at 37 °C in fresh media to wash off unbound substrate. Imaging was performed using FluoroBrite DMEM (Gibco) supplemented with 10% fetal calf serum.

FRAP experiments were carried out on a Leica Stellaris 8 using a 20mW 488nm Diode laser and a 63x 1.4NA oil immersion lens. Images were acquired every 2 s using the Resonant Scanner with line averaging of 2. Pixel size was set to nyquist (103 nm) sampling and 256 x 256 array was acquired. Bleaching parameters: in all image series 5 ‘pre-bleach’ images were acquired, a consistently sized area of 2 µm diameter was bleached. Bleach settings were 488 nm laser set to 100% 15 repetitions. Post-bleach images were acquired every 2 s for 150 s. FRAP analysis was performed using custom scripts, following a protocol similar to that described by Bancaud et al., (Bancaud et al., 2010). Cell movements were corrected by registering (through roto-traslations) all frames with the starting one using the TurboReg plugin in Fiji (Thevenaz et al., 1998). The mean intensity of the bleaching region *I^B^*(*t*), as well as that of the entire nucleus *I^C^*(*t*) was measured. A first normalisation was performed to correct the photobleaching of the signal in time and to normalise the signal in the range [0,1], where 1 corresponds to the normalised pre-bleaching signal:

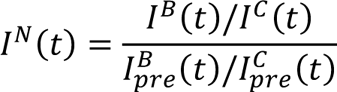

where 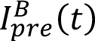 and 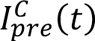 are respectively the average pre-bleach signal in the bleached region and in the entire nucleus.

A second normalisation was performed to shift the minimum intensity of the signal to 0, while maintaining the signal in the range of 0 to 1. This is necessary for visually comparing FRAP curves for different conditions, as they typically have different minima in the intensity (bleaching is not 100% efficient). This second normalisation does not influence the fitting of the curves. The new normalised intensity is therefore:

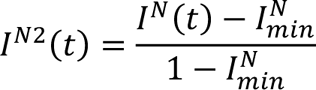

Being 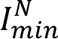 the minimum intensity of the FRAP curve.

For every FRAP recovery curve, we calculated the fraction of the immobile (or very slow) population, defined as *IF* 1 – *I*^*N*2^(*t* = +∞). In our case we approximated *I*^*N*2^(*t* = +∞) as the average of the last 50 values of *I*^*N*2^ (i.e., from *t* = 100*s* to *t* = 150*s*). The FRAP recovery curves and immobile fraction were calculated for every experiment, as well as their standard deviation. Data shown in *Fig. 6e* are obtained from the mean of the 3 biological repeats (experiment 1: WT n=3, ΔN n=6; experiment 2: WT n=5, ΔN n=15; experiment 3: WT n=10, ΔN n=7). Standard deviation has been calculated using the theory of errors propagation for uncorrelated data.

## Code availability

Custom scripts used in the analysis of data are available from the authors upon request.

## Data availability

All sequencing data that were generated during this study will be deposited in GEO and made available upon publication.

## Acknowledgements

We thank C. Uggenti, P. Heyn, A. P. Jackson, Y. Crow, S. Janssen, M. Lorincz and members of the Sproul lab for useful discussions. We thank Edinburgh Clinical Research Facility Genetics Core, MRC IGC FACs and imaging core facilities for technical support. This work has made use of the resources provided by the University of Edinburgh digital research services and the MRC IGMM compute cluster. D.S. is a Cancer Research UK Career Development fellow (reference C47648/A20837), and work in his laboratory is also supported by an MRC university grant to the MRC Human Genetics Unit. M.D.W.’s work is supported by the Wellcome Trust (210493), Medical Research Council (T029471/1), and the University of Edinburgh. This work was supported by the Edinburgh Protein Production Facility (EPPF), which receives funding from a core grant (203149) to the Wellcome Centre for Cell Biology at the University of Edinburgh. LK is a cross-disciplinary post-doctoral fellow supported by funding from the University of Edinburgh and Medical Research Council (MC\_UU\_00009/2). I.K. was funded by a studentship from CRUK as well as the A. G. Leventis Foundation. H.F. was funded by an ERASMUS+ scholarship.

## Contributions

F.T., K.I.M., I.K., H.Y.L, Y.Z., M.M., H.F., C.R.R., H.W., H.D.S., and M.D.W. performed the experiments included in the manuscript. J.W. conducted mass spectrometry. A.W. supervised the FRAP experiment. L.C.M assisted with image analysis. F.T., I.K., M.M. L.K. and D.S. conducted the computational analysis presented in the manuscript. D.S., F.T. and M.D.W. planned and supervised the study. D.S. and F.T. wrote the manuscript with input from and review by all authors.

## References

Allshire, R.C., and Madhani, H.D. (2018). Ten principles of heterochromatin formation and function. Nat Rev Mol Cell Biol 19, 229–244.

Amemiya, H.M., Kundaje, A., and Boyle, A.P. (2019). The ENCODE Blacklist: Identification of Problematic Regions of the Genome. Sci Rep 9, 9354.

Ashkenazy, H., Abadi, S., Martz, E., Chay, O., Mayrose, I., Pupko, T., and Ben-Tal, N. (2016). ConSurf 2016: an improved methodology to estimate and visualize evolutionary conservation in macromolecules. Nucleic Acids Res 44, W344–350.

Bachman, K.E., Rountree, M.R., and Baylin, S.B. (2001). Dnmt3a and Dnmt3b are transcriptional repressors that exhibit unique localization properties to heterochromatin. J Biol Chem 276, 32282–32287.

Bancaud, A., Huet, S., Rabut, G., and Ellenberg, J. (2010). Fluorescence perturbation techniques to study mobility and molecular dynamics of proteins in live cells: FRAP, photoactivation, photoconversion, and FLIP. Cold Spring Harb Protoc 2010, pdb top90.

Bannister, A.J., Schneider, R., Myers, F.A., Thorne, A.W., Crane-Robinson, C., and Kouzarides, T. (2005). Spatial distribution of di- and tri-methyl lysine 36 of histone H3 at active genes. J Biol Chem 280, 17732–17736.

Baubec, T., Colombo, D.F., Wirbelauer, C., Schmidt, J., Burger, L., Krebs, A.R., Akalin, A., and Schübeler, D. (2015). Genomic profiling of DNA methyltransferases reveals a role for DNMT3B in genic methylation. Nature 520, 243–247.

Blackledge, N.P., Rose, N.R., and Klose, R.J. (2015). Targeting Polycomb systems to regulate gene expression: modifications to a complex story. Nat Rev Mol Cell Biol 16, 643–649.

Boyko, K., Arkova, O., Nikolaeva, A., Popov, V.O., Georgiev, P., and Bonchuk, A. (2022). Structure of the DNMT3B ADD domain suggests the absence of a DNMT3A-like autoinhibitory mechanism. Biochem Biophys Res Commun 619, 124–129.

Brandle, F., Fruhbauer, B., and Jagannathan, M. (2022). Principles and functions of pericentromeric satellite DNA clustering into chromocenters. Semin Cell Dev Biol 128, 26–39.

Chen, T., Tsujimoto, N., and Li, E. (2004). The PWWP domain of Dnmt3a and Dnmt3b is required for directing DNA methylation to the major satellite repeats at pericentric heterochromatin. Mol Cell Biol 24, 9048–9058.

Chen, T., Ueda, Y., Xie, S., and Li, E. (2002). A novel Dnmt3a isoform produced from an alternative promoter localizes to euchromatin and its expression correlates with active de novo methylation. J Biol Chem 277, 38746–38754.

Chen, Z.X., Mann, J.R., Hsieh, C.L., Riggs, A.D., and Chédin, F. (2005). Physical and functional interactions between the human DNMT3L protein and members of the de novo methyltransferase family. Journal of Cellular Biochemistry 95, 902–917.

Decato, B.E., Qu, J., Ji, X., Wagenblast, E., Knott, S.R.V., Hannon, G.J., and Smith, A.D. (2020). Characterization of universal features of partially methylated domains across tissues and species. Epigenetics Chromatin 13, 39.

Dhayalan, A., Rajavelu, A., Rathert, P., Tamas, R., Jurkowska, R.Z., Ragozin, S., and Jeltsch, A. (2010). The Dnmt3a PWWP domain reads histone 3 lysine 36 trimethylation and guides DNA methylation. Journal of Biological Chemistry 285, 26114–26120.

Eidahl, J.O., Crowe, B.L., North, J.A., McKee, C.J., Shkriabai, N., Feng, L., Plumb, M., Graham, R.L., Gorelick, R.J., Hess, S., et al. (2013). Structural basis for high-affinity binding of LEDGF PWWP to mononucleosomes. Nucleic Acids Res 41, 3924–3936.

Elliott, E.N., Sheaffer, K.L., and Kaestner, K.H. (2016). The ‘de novo’ DNA methyltransferase Dnmt3b compensates the Dnmt1-deficient intestinal epithelium. eLife 5, 1–15.

Gao, L., Guo, Y., Biswal, M., Lu, J., Yin, J., Fang, J., Chen, X., Shao, Z., Huang, M., Wang, Y., et al. (2022). Structure of DNMT3B homo-oligomer reveals vulnerability to impairment by ICF mutations. Nat Commun 13, 4249.

Gatto, S., Gagliardi, M., Franzese, M., Leppert, S., Papa, M., Cammisa, M., Grillo, G., Velasco, G., Francastel, C., Toubiana, S., et al. (2017). ICF-specific DNMT3B dysfunction interferes with intragenic regulation of mRNA transcription and alternative splicing. Nucleic Acids Research 45, 5739–5756.

Ge, Y.Z., Pu, M.T., Gowher, H., Wu, H.P., Ding, J.P., Jeltsch, A., and Xu, G.L. (2004). Chromatin targeting of de novo DNA methyltransferases by the PWWP domain. Journal of Biological Chemistry 279, 25447–25454.

Goll, M.G., and Bestor, T.H. (2005). Eukaryotic cytosine methyltransferases. Annu Rev Biochem 74, 481–514.

Gu, T., Hao, D., Woo, J., Huang, T.W., Guo, L., Lin, X., Guzman, A.G., Tovy, A., Rosas, C., Jeong, M., et al. (2022). The disordered N-terminal domain of DNMT3A recognizes H2AK119ub and is required for postnatal development. Nat Genet 54, 625–636.

Guo, X., Wang, L., Li, J., Ding, Z., Xiao, J., Yin, X., He, S., Shi, P., Dong, L., Li, G., et al. (2015). Structural insight into autoinhibition and histone H3-induced activation of DNMT3A. Nature 517, 640–644.

Hassan, K.M., Norwood, T., Gimelli, G., Gartler, S.M., and Hansen, R.S. (2001). Satellite 2 methylation patterns in normal and ICF syndrome cells and association of hypomethylation with advanced replication. Hum Genet 109, 452–462.

Heyn, H., Vidal, E., Sayols, S., Sanchez-Mut, J.V., Moran, S., Medina, I., Sandoval, J., Simo-Riudalbas, L., Szczesna, K., Huertas, D., et al. (2012). Whole-genome bisulfite DNA sequencing of a DNMT3B mutant patient. Epigenetics 7, 542–550.

Heyn, P., Logan, C.V., Fluteau, A., Challis, R.C., Auchynnikava, T., Martin, C.A., Marsh, J.A., Taglini, F., Kilanowski, F., Parry, D.A., et al. (2019). Gain-of-function DNMT3A mutations cause microcephalic dwarfism and hypermethylation of Polycomb-regulated regions. Nature Genetics 51, 96–105.

Hsieh, C.L. (1999). In vivo activity of murine de novo methyltransferases, Dnmt3a and Dnmt3b. Mol Cell Biol 19, 8211–8218.

Hu, J.L., Zhou, B.O., Zhang, R.R., Zhang, K.L., Zhou, J.Q., and Xu, G.L. (2009). The N-terminus of histone H3 is required for de novo DNA methylation in chromatin. Proc Natl Acad Sci U S A 106, 22187–22192.

Huang, Y.H., Chen, C.W., Sundaramurthy, V., Slabicki, M., Hao, D., Watson, C.J., Tovy, A., Reyes, J.M., Dakhova, O., Crovetti, B.R., et al. (2022). Systematic Profiling of DNMT3A Variants Reveals Protein Instability Mediated by the DCAF8 E3 Ubiquitin Ligase Adaptor. Cancer Discov 12, 220–235.

Jackson, M.S., Mole, S.E., and Ponder, B.A. (1992). Characterisation of a boundary between satellite III and alphoid sequences on human chromosome 10. Nucleic Acids Res 20, 4781–4787.

Jeanpierre, M., Turleau, C., Aurias, A., Prieur, M., Ledeist, F., Fischer, A., and Viegas-Pequignot, E. (1993). An embryonic-like methylation pattern of classical satellite DNA is observed in ICF syndrome. Hum Mol Genet 2, 731–735.

Jeltsch, A., and Jurkowska, R.Z. (2016). Allosteric control of mammalian DNA methyltransferases - a new regulatory paradigm. Nucleic Acids Res 44, 8556–8575.

Jeong, S., Liang, G., Sharma, S., Lin, J.C., Choi, S.H., Han, H., Yoo, C.B., Egger, G., Yang, A.S., and Jones, P.A. (2009). Selective Anchoring of DNA Methyltransferases 3A and 3B to Nucleosomes Containing Methylated DNA. Molecular and Cellular Biology 29, 5366–5376.

Krueger, F., and Andrews, S.R. (2011). Bismark: a flexible aligner and methylation caller for Bisulfite-Seq applications. Bioinformatics 27, 1571–1572.

Langmead, B., and Salzberg, S.L. (2012). Fast gapped-read alignment with Bowtie 2. Nat Methods 9, 357–359.

Lehnertz, B., Ueda, Y., Derijck, A.A., Braunschweig, U., Perez-Burgos, L., Kubicek, S., Chen, T., Li, E., Jenuwein, T., and Peters, A.H. (2003). Suv39h-mediated histone H3 lysine 9 methylation directs DNA methylation to major satellite repeats at pericentric heterochromatin. Curr Biol 13, 1192–1200.

Li, B.Z., Huang, Z., Cui, Q.Y., Song, X.H., Du, L., Jeltsch, A., Chen, P., Li, G., Li, E., and Xu, G.L. (2011). Histone tails regulate DNA methylation by allosterically activating de novo methyltransferase. Cell Res 21, 1172–1181.

Li, H., Handsaker, B., Wysoker, A., Fennell, T., Ruan, J., Homer, N., Marth, G., Abecasis, G., Durbin, R., and Genome Project Data Processing, S. (2009). The Sequence Alignment/Map format and SAMtools. Bioinformatics 25, 2078–2079.

Liang, G., Chan, M.F., Tomigahara, Y., Tsai, Y.C., Gonzales, F.A., Li, E., Laird, P.W., and Jones, P.A. (2002). Cooperativity between DNA methyltransferases in the maintenance methylation of repetitive elements. Mol Cell Biol 22, 480–491.

Liao, J., Karnik, R., Gu, H., Ziller, M.J., Clement, K., Tsankov, A.M., Akopian, V., Gifford, C.A., Donaghey, J., Galonska, C., et al. (2015). Targeted disruption of DNMT1, DNMT3A and DNMT3B in human embryonic stem cells. Nature Genetics 47, 469–478.

Lue, N.Z., Garcia, E.M., Ngan, K.C., Lee, C., Doench, J.G., and Liau, B.B. (2022). Base editor scanning charts the DNMT3A activity landscape. bioRxiv, 2022.2004.2012.487946.

Macdonald, J., Taylor, L., Sherman, A., Kawakami, K., Takahashi, Y., Sang, H.M., and McGrew, M.J. (2012). Efficient genetic modification and germ-line transmission of primordial germ cells using piggyBac and Tol2 transposons. Proc Natl Acad Sci U S A 109, E1466–1472.

Manzo, M., Wirz, J., Ambrosi, C., Villasenor, R., Roschitzki, B., and Baubec, T. (2017). Isoform-specific localization of DNMT3A regulates DNA methylation fidelity at bivalent CpG islands. EMBO J 36, 3421–3434.

Masalmeh, R.H.A., Taglini, F., Rubio-Ramon, C., Musialik, K.I., Higham, J., Davidson-Smith, H., Kafetzopoulos, I., Pawlicka, K.P., Finan, H.M., Clark, R., et al. (2021). De novo DNA methyltransferase activity in colorectal cancer is directed towards H3K36me3 marked CpG islands. Nat Commun 12, 694.

Morselli, M., Pastor, W.A., Montanini, B., Nee, K., Ferrari, R., Fu, K., Bonora, G., Rubbi, L., Clark, A.T., Ottonello, S., et al. (2015). In vivo targeting of de novo DNA methylation by histone modifications in yeast and mouse. eLife 2015, 1–21.

Neri, F., Rapelli, S., Krepelova, A., Incarnato, D., Parlato, C., Basile, G., Maldotti, M., Anselmi, F., and Oliviero, S. (2017). Intragenic DNA methylation prevents spurious transcription initiation. Nature 543, 72–77.

Okano, M., Bell, D.W., Haber, D.A., and Li, E. (1999). DNA Methyltransferases Dnmt3a and Dnmt3b Are Essential for De Novo Methylation and Mammalian Development. Cell 99, 247–257.

Orlando, D.A., Chen, M.W., Brown, V.E., Solanki, S., Choi, Y.J., Olson, E.R., Fritz, C.C., Bradner, J.E., and Guenther, M.G. (2014). Quantitative ChIP-Seq normalization reveals global modulation of the epigenome. Cell Rep 9, 1163–1170.

Otani, J., Nankumo, T., Arita, K., Inamoto, S., Ariyoshi, M., and Shirakawa, M. (2009). Structural basis for recognition of H3K4 methylation status by the DNA methyltransferase 3A ATRX-DNMT3-DNMT3L domain. EMBO Reports 10, 1235–1241.

Pchelintsev, N.A., Adams, P.D., and Nelson, D.M. (2016). Critical Parameters for Efficient Sonication and Improved Chromatin Immunoprecipitation of High Molecular Weight Proteins. PLoS One 11, e0148023.

Pettersen, E.F., Goddard, T.D., Huang, C.C., Couch, G.S., Greenblatt, D.M., Meng, E.C., and Ferrin, T.E. (2004). UCSF Chimera--a visualization system for exploratory research and analysis. J Comput Chem 25, 1605–1612.

Quinlan, A.R., and Hall, I.M. (2010). BEDTools: a flexible suite of utilities for comparing genomic features. Bioinformatics 26, 841–842.

Ramirez, F., Ryan, D.P., Gruning, B., Bhardwaj, V., Kilpert, F., Richter, A.S., Heyne, S., Dundar, F., and Manke, T. (2016). deepTools2: a next generation web server for deep-sequencing data analysis. Nucleic Acids Res 44, W160–165.

Rhee, I., Bachman, K.E., Park, B.H., Jair, K.-W., Yen, R.-W.C., Schuebel, K.E., Cui, H., Feinberg, A.P., Lengauer, C., Kinzler, K.W., et al. (2002). DNMT1 and DNMT3b cooperate to silence genes in human cancer cells. Nature 416, 552–556.

Rondelet, G., Dal Maso, T., Willems, L., and Wouters, J. (2016). Structural basis for recognition of histone H3K36me3 nucleosome by human de novo DNA methyltransferases 3A and 3B. Journal of Structural Biology 194, 357–367.

Sendžikaitė, G., Hanna, C.W., Stewart-Morgan, K.R., Ivanova, E., and Kelsey, G. (2019). A DNMT3A PWWP mutation leads to methylation of bivalent chromatin and growth retardation in mice. Nature Communications 10.

Shirohzu, H., Kubota, T., Kumazawa, A., Sado, T., Chijiwa, T., Inagaki, K., Suetake, I., Tajima, S., Wakui, K., Miki, Y., et al. (2002). Three novel DNMT3B mutations in Japanese patients with ICF syndrome. American Journal of Medical Genetics 112, 31–37.

Smith, C.A., O’Maille, G., Want, E.J., Qin, C., Trauger, S.A., Brandon, T.R., Custodio, D.E., Abagyan, R., and Siuzdak, G. (2005). METLIN: a metabolite mass spectral database. Ther Drug Monit 27, 747–751.

Smith, T., Heger, A., and Sudbery, I. (2017). UMI-tools: modeling sequencing errors in Unique Molecular Identifiers to improve quantification accuracy. Genome Res 27, 491–499.

Spracklin, G., and Pradhan, S. (2020). Protect-seq: genome-wide profiling of nuclease inaccessible domains reveals physical properties of chromatin. Nucleic Acids Res 48, e16.

Suzuki, M.M., and Bird, A. (2008). DNA methylation landscapes: provocative insights from epigenomics. Nat Rev Genet 9, 465–476.

Tarasov, A., Vilella, A.J., Cuppen, E., Nijman, I.J., and Prins, P. (2015). Sambamba: fast processing of NGS alignment formats. Bioinformatics 31, 2032–2034.

Thevenaz, P., Ruttimann, U.E., and Unser, M. (1998). A pyramid approach to subpixel registration based on intensity. IEEE Trans Image Process 7, 27–41.

Tian, W., Yan, P., Xu, N., Chakravorty, A., Liefke, R., Xi, Q., and Wang, Z. (2019). The HRP3 PWWP domain recognizes the minor groove of double-stranded DNA and recruits HRP3 to chromatin. Nucleic Acids Res 47, 5436–5448.

van Nuland, R., van Schaik, F.M., Simonis, M., van Heesch, S., Cuppen, E., Boelens, R., Timmers, H.M., and van Ingen, H. (2013). Nucleosomal DNA binding drives the recognition of H3K36-methylated nucleosomes by the PSIP1-PWWP domain. Epigenetics Chromatin 6, 12.

van Steensel, B., and Belmont, A.S. (2017). Lamina-Associated Domains: Links with Chromosome Architecture, Heterochromatin, and Gene Repression. Cell 169, 780–791.

Wang, H., Farnung, L., Dienemann, C., and Cramer, P. (2020). Structure of H3K36-methylated nucleosome-PWWP complex reveals multivalent cross-gyre binding. Nat Struct Mol Biol 27, 8–13.

Waterhouse, A.M., Procter, J.B., Martin, D.M., Clamp, M., and Barton, G.J. (2009). Jalview Version 2-- a multiple sequence alignment editor and analysis workbench. Bioinformatics 25, 1189–1191.

Weemaes, C.M., van Tol, M.J., Wang, J., van Ostaijen-ten Dam, M.M., van Eggermond, M.C., Thijssen, P.E., Aytekin, C., Brunetti-Pierri, N., van der Burg, M., Graham Davies, E., et al. (2013). Heterogeneous clinical presentation in ICF syndrome: correlation with underlying gene defects. Eur J Hum Genet 21, 1219–1225.

Weinberg, D.N., Papillon-Cavanagh, S., Chen, H., Yue, Y., Chen, X., Rajagopalan, K.N., Horth, C., McGuire, J.T., Xu, X., Nikbakht, H., et al. (2019). The histone mark H3K36me2 recruits DNMT3A and shapes the intergenic DNA methylation landscape. Nature 573, 281–286.

Weinberg, D.N., Rosenbaum, P., Chen, X., Barrows, D., Horth, C., Marunde, M.R., Popova, I.K., Gillespie, Z.B., Keogh, M.C., Lu, C., et al. (2021). Two competing mechanisms of DNMT3A recruitment regulate the dynamics of de novo DNA methylation at PRC1-targeted CpG islands. Nat Genet 53, 794–800.

Weisenberger, D.J., Velicescu, M., Cheng, J.C., Gonzales, F.a., Liang, G., and Jones, P.a. (2004). Role of the DNA methyltransferase variant DNMT3b3 in DNA methylation. Molecular cancer research : MCR 2, 62–72.

Wilson, M.D., Renault, L., Maskell, D.P., Ghoneim, M., Pye, V.E., Nans, A., Rueda, D.S., Cherepanov, P., and Costa, A. (2019). Retroviral integration into nucleosomes through DNA looping and sliding along the histone octamer. Nat Commun 10, 4189.

Xu, G.L., Bestor, T.H., Bourc’his, D., Hsieh, C.L., Tommerup, N., Bugge, M., Hulten, M., Qu, X., Russo, J.J., and Viegas-Pequignot, E. (1999). Chromosome instability and immunodeficiency syndrome caused by mutations in a DNA methyltransferase gene. Nature 402, 187–191.

Zhang, M., Lei, M., Qin, S., Dong, A., Yang, A., Li, Y., Loppnau, P., Hughes, T.R., Min, J., and Liu, Y. (2021). Crystal structure of the BRPF2 PWWP domain in complex with DNA reveals a different binding mode than the HDGF family of PWWP domains. Biochim Biophys Acta Gene Regul Mech 1864, 194688.

Zhang, Y., Jurkowska, R., Soeroes, S., Rajavelu, A., Dhayalan, A., Bock, I., Rathert, P., Brandt, O., Reinhardt, R., Fischle, W., et al. (2010). Chromatin methylation activity of Dnmt3a and Dnmt3a/3L is guided by interaction of the ADD domain with the histone H3 tail. Nucleic Acids Research 38, 4246–4253.

Zhou, W., Dinh, H.Q., Ramjan, Z., Weisenberger, D.J., Nicolet, C.M., Shen, H., Laird, P.W., and Berman, B.P. (2018). DNA methylation loss in late-replicating domains is linked to mitotic cell division. Nat Genet 50, 591–602.

