## Supplementary materials for "DNMT3B PWWP mutations cause hypermethylation of heterochromatin"

#### Supplementary Table 1

*Oligonucleotides used in this study.*

| CRISPR/Cas9-edited HCT116 cell line | |
|---|---|
| **Name** | **Sequence** |
| DNMT3B-EX7-W263A-HDR-Template | CAAGCGACAGGCTATGTCTGGCATGCGATGGGTCCAGGCGTTTGGCGATGGCAAGTTCTCCGAGGTGAGTCCGGGGAAGG |
| gRNA-EX7-T | [Phos]CACCGACTTGCCATCGCCAAACCAC |
| gRNA-EX7-B | [Phos]AAACGTGGTTTGGCGATGGCAAGTC |
| DNMT3B-EX7-screen-F | TTTGGAATAGGGGACCTCGTGTGG |
| DNMT3B-EX7-screen-R | CACACATCTGCAGAATACAATCCCAGG |
| **bsPCR primers** | |
| **Name** | **Sequence** |
| bs-BRCA2-UMI-3-F | TCGTCGGCAGCGTCAGATGTGTATAAGAGACAGNNNNGTTTGGGATTTTTAAGGGTTAG |
| bs-BRCA2-UMI-3-R | GTCTCGTGGGCTCGGAGATGTGTATAAGAGACNNNNCTCCAAAAATACAATTCTTTTT |
| bsH3K9me3L1-UMI-F | TCGTCGGCAGCGTCAGATGTGTATAAGAGACAGNNNNGAGTTTAGTATTTTTTTTCGGAGTT |
| bsH3K9me3L1-UMI-R | GTCTCGTGGGCTCGGAGATGTGTATAAGAGACNNNNCCCTCCTCCTTAAAATAAAAAAAAC |
| bsH3K9me3L2-UMI-F | TCGTCGGCAGCGTCAGATGTGTATAAGAGACAGNNNNGAAGAGGAGGGTGGTTGGGAAGGGTG |
| bsH3K9me3L2-UMI-R | GTCTCGTGGGCTCGGAGATGTGTATAAGAGACNNNNCATAATCACATAAACAAAAACCACAA |

#### Supplementary Table 2

*Summary of sequencing statistics for WGBS. Aligned reads counts are following PCR duplicate removal.*

| Sample | Total reads ($x10^6$) | Aligned reads ($x10^6$) | Mean CG coverage | Bisulfite conversion rate |
|---|---|---|---|---|
| 3BKO vs HCT116 and expression of 3B and 3Bcd in 3BKO cells - Figure 1 and S1 | | | | |
| HCT116 | 43.40 | 32.72 | 2.53 | 99.28 |
| 3BKO | 52.38 | 39.73 | 2.75 | 99.28 |
| 3BKO+3B | 61.34 | 47.06 | 2.66 | 99.28 |
| 3BKO+3Bcd | 48.45 | 36.89 | 2.53 | 99.28 |
| expression of 3A and 3B W263A in 3BKO cells - Figure 2 and S2 | | | | |
| HCT116 | 84.54 | 64.46 | 2.55 | 99.70 |
| 3BKO | 116.48 | 90.62 | 3.11 | 99.69 |
| 3BKO+3B | 94.42 | 72.72 | 2.67 | 99.68 |
| 3BKO+3BW263A | 91.77 | 69.95 | 2.57 | 99.69 |
| 3BKO+3A | 116.63 | 90.44 | 3.02 | 99.67 |
| 3BKO+3Acd | 92.15 | 70.65 | 2.54 | 99.57 |
| control and 3B W263A CRISPR clones - Figure 2 and S2 | | | | |
| HCT116 | 44.07 | 43.03 | 2.02 | 99.64 |
| W263A c1 | 58.61 | 57.72 | 2.43 | 99.68 |
| W263A c2 | 60.38 | 59.64 | 2.50 | 99.70 |
| W263A c3 | 70.46 | 69.78 | 2.74 | 99.66 |
| control c1 | 87.63 | 86.98 | 3.18 | 99.67 |
| control c2 | 69.60 | 68.79 | 2.68 | 99.66 |
| expression of 3B ΔN in 3BKO cells- Figure 6 and S6 | | | | |
| HCT116 | 45.08 | 35.72 | 1.88 | 99.68 |
| 3BKO | 49.79 | 39.44 | 2.01 | 99.64 |
| 3BKO+3B rep.1 | 46.77 | 36.77 | 1.97 | 99.52 |
| 3BKO+3BΔN rep.1 | 47.28 | 37.05 | 1.99 | 99.58 |
| 3BKO+3B rep.2 | 50.25 | 39.81 | 2.01 | 99.60 |
| 3BKO+3BΔN rep.2 | 46.77 | 36.90 | 1.98 | 99.57 |

**Supplementary Table 3**

*Summary of sequencing statistics for ChIP-Rx-seq. Aligned reads are after duplicate and multi- mapper removal and reported for both genomes (hg38/dm6). For T7-DNMT3B ChIP-Rx-seq, which was paired end sequencing, fragments are quoted.*

| Sample | Total reads or read pairs (x$10^6$) | Aligned reads or fragments hg38 (x$10^6$) or hg38 (x$10^6$)/dm6 (x$10^3$) |
|---|---|---|
| | H3K9me3 and H3K27me3 | |
| HCT116 Input rep. 1 | 41.49 | 32.98 |
| DNMT3B KO Input rep. 1 | 55.10 | 43.87 |
| HCT116 H3K9me3 rep. 1 | 37.95 | 29.60 |
| DNMT3B KO H3K9me3 rep. 1 | 34.10 | 26.71 |
| HCT116 H3K27me3 rep. 1 | 52.57 | 40.43 |
| DNMT3B KO H3K27me3 rep. 1 | 35.81 | 28.57 |
| HCT116 Input rep. 2 | 50.59 | 39.79 |
| DNMT3B KO Input rep. 2 | 53.15 | 42.16 |
| HCT116 H3K9me3 rep. 2 | 29.01 | 22.61 |
| DNMT3B KO H3K9me3 rep. 2 | 45.66 | 35.86 |
| HCT116 H3K27me3 rep. 2 | 51.08 | 40.08 |
| DNMT3B KO H3K27me3 rep. 2 | 60.89 | 48.65 |
| | H3K36me3 | |
| HCT116 Input rep. 1 | 70.54 | 59.19/110 |
| HCT116 H3K36me3 rep. 1 | 70.07 | 58.09/290 |
| DNMT3B KO Input rep. 1 | 63.62 | 53.71/100 |
| DNMT3B KO H3K36me3 rep. 1 | 66.01 | 54.92/250 |
| HCT116 Input rep. 2 | 72.15 | 60.51/130 |
| HCT116 H3K36me3 rep. 2 | 70.96 | 58.78/280 |
| DNMT3B KO Input rep. 2 | 67.03 | 56.62/110 |
| DNMT3B KO H3K36me3 rep. 2 | 67.84 | 56.57/220 |
| | H3K4me3 | |
| HCT116 Input rep. 1 | 107.75 | 84.71 |
| DNMT3B KO Input rep. 1 | 74.93 | 59.03 |
| HCT116 H3K4me3 rep. 1 | 83.08 | 64.17 |
| DNMT3B KO H3K4me3 rep. 1 | 51.52 | 40.69 |
| HCT116 Input rep. 2 | 58.10 | 45.77 |
| DNMT3B KO Input rep. 2 | 33.20 | 25.93 |
| HCT116 H3K4me3 rep. 2 | 95.11 | 72.07 |
| DNMT3B KO H3K4me3 rep. 2 | 79.37 | 60.17 |
| | endogenous T7-DNMT3B | |
| HCT116 (mock) Input rep. 1 | 74.80 | 63.42/5 |
| HCT116 (mock) T7-IP rep. 1 | 52.35 | 39.36/1902 |
| T7-DNMT3B Input rep. 1 | 78.31 | 66.45/5 |
| T7-DNMT3B T7-IP rep. 1 | 73.32 | 62.90/332 |
| HCT116 (mock) Input rep. 2 | 74.94 | 63.58/6 |
| HCT116 T7-IP (mock) rep. 2 | 66.10 | 56.25/85 |
| T7-DNMT3B Input rep. 2 | 68.89 | 57.81/6 |
| T7-DNMT3B T7-IP rep. 2 | 83.90 | 70.50/179 |
| | T7-DNMT3B in WT cells | |
| HCT116 (mock) Input rep. 1 | 34.21 | 25.62 |
| HCT116 +T7-DNMT3B Input rep. 1 | 50.94 | 37.63 |
| HCT116 (mock) T7-IP rep. 1 | 32.82 | 24.67 |
| HCT116 +T7-DNMT3B T7-IP rep. 1 | 52.42 | 39.48 |
| HCT116 (mock) Input rep. 2 | 66.06 | 47.61 |
| HCT116 +T7-DNMT3B Input rep. 2 | 39.87 | 29.26 |
| HCT116 (mock) T7-IP rep. 2 | 28.86 | 21.65 |
| HCT116 +T7-DNMT3B T7-IP rep. 2 | 37.13 | 28.05 |
| | T7-DNMT3B in 3BKO cells | |
| 3BKO (mock) Input rep. 1 | 69.38 | 49.07/3.15 |
| 3BKO +T7-DNMT3B Input rep. 1 | 89.62 | 61.21/4.93 |
| 3BKO +T7-DNMT3BW263A Input rep. 1 | 108.42 | 73.43/4.36 |
| 3BKO +T7-DNMT3BΔN Input rep. 1 | 92.92 | 64.51/2.23 |
| 3BKO (mock) T7-IP rep. 1 | 53.30 | 38.06/33.96 |
| 3BKO +T7-DNMT3B T7-IP rep. 1 | 81.88 | 57.86/10.79 |
| 3BKO +T7-DNMT3BW263A T7-IP rep. 1 | 76.48 | 53.87/14.16 |
| 3BKO +T7-DNMT3BΔN T7-IP rep. 1 | 81.00 | 58.17/6.33 |

| | | |
|---|---|---|
| 3BKO (mock) Input rep. 2 | 71.84 | 49.91/3.28 |
| 3BKO +T7-DNMT3B Input rep. 2 | 94.73 | 65.36/2.32 |
| 3BKO +T7-DNMT3BW263A Input rep. 2 | 89.59 | 62.45/1.60 |
| 3BKO +T7-DNMT3BΔN Input rep. 2 | 87.19 | 61.60/1.52 |
| 3BKO (mock) T7-IP rep. 2 | 65.52 | 45.72/34.06 |
| 3BKO +T7-DNMT3B T7-IP rep. 2 | 77.87 | 56.56/8.73 |
| 3BKO +T7-DNMT3BW263A T7-IP rep. 2 | 73.14 | 52.74/9.73 |
| 3BKO +T7-DNMT3BΔN T7-IP rep. 2 | 73.46 | 53.63/5.67 |

**Supplementary Table 4**

*Summary of bisulfite PCR reads following PCR duplicate removal.*

| Sample | Locus | Number of reads |
|---|---|---|
| D266A experiment | | |
| 3BKO | BRCA2 | 15 |
| 3BKO+3B | | 10 |
| 3BKO+3B W263A | | 7 |
| 3BKO+3B D266A | | 16 |
| 3BKO+eGFP | | 11 |
| 3BKO | H3K9me3 locus 1 | 56 |
| 3BKO+3B | | 43 |
| 3BKO+3B W263A | | 44 |
| 3BKO+3B D266A | | 56 |
| 3BKO+eGFP | | 24 |
| 3BKO | H3K9me3 locus 2 | 156 |
| 3BKO+3B | | 148 |
| 3BKO+3B W263A | | 166 |
| 3BKO+3B D266A | | 196 |
| 3BKO+eGFP | | 162 |
| S270P experiment | | |
| 3BKO | BRCA2 | 22 |
| 3BKO+3B | | 22 |
| 3BKO+3B W263A | | 7 |
| 3BKO+3B S270P | | 13 |
| 3BKO+eGFP | | 12 |
| 3BKO | H3K9me3 locus 1 | 84 |
| 3BKO+3B | | 75 |
| 3BKO+3B W263A | | 22 |
| 3BKO+3B S270P | | 105 |
| 3BKO+eGFP | | 43 |
| 3BKO | H3K9me3 locus 2 | 140 |
| 3BKO+3B | | 145 |
| 3BKO+3B W263A | | 155 |
| 3BKO+3B S270P | | 141 |
| 3BKO+eGFP | | 202 |
| PWWP and ΔN mutants experiment | | |
| 3BKO | BRCA2 | 101 |
| 3BKO+3B | | 55 |
| 3BKO+3B W263A | | 38 |
| 3BKO+3B K276E | | 77 |
| 3BKO+3B K294E | | 64 |
| 3BKO+3B W263A K276E | | 80 |
| 3BKO+3B W263A K294E | | 74 |
| 3BKO+3B ΔPWWP | | 75 |
| 3BKO+3B ΔN | | 66 |
| 3BKO+3B ΔN W263A | | 100 |
| 3BKO+eGFP | | 46 |
| 3BKO | H3K9me3 locus 1 | 1652 |
| 3BKO+3B | | 797 |
| 3BKO+3B W263A | | 1078 |
| 3BKO+3B K276E | | 1449 |
| 3BKO+3B K294E | | 695 |
| 3BKO+3B W263A K276E | | 1244 |
| 3BKO+3B W263A K294E | | 863 |
| 3BKO+3B ΔPWWP | | 605 |
| 3BKO+3B ΔN | | 1198 |
| 3BKO+3B ΔN W263A | | 1025 |
| 3BKO+eGFP | | 791 |
| 3BKO | H3K9me3 locus 2 | 5363 |
| 3BKO+3B | | 4772 |
| 3BKO+3B W263A | | 4332 |
| 3BKO+3B K276E | | 5161 |
| 3BKO+3B K294E | | 4157 |
| 3BKO+3B W263A K276E | | 5249 |
| 3BKO+3B W263A K294E | | 4824 |

| | | |
|---|---|---|
| 3BKO+3B ΔPWWP | | 4979 |
| 3BKO+3B ΔN | | 5191 |
| 3BKO+3B ΔN W263A | | 5135 |
| 3BKO+eGFP | | 4035 |

### Fig. S1

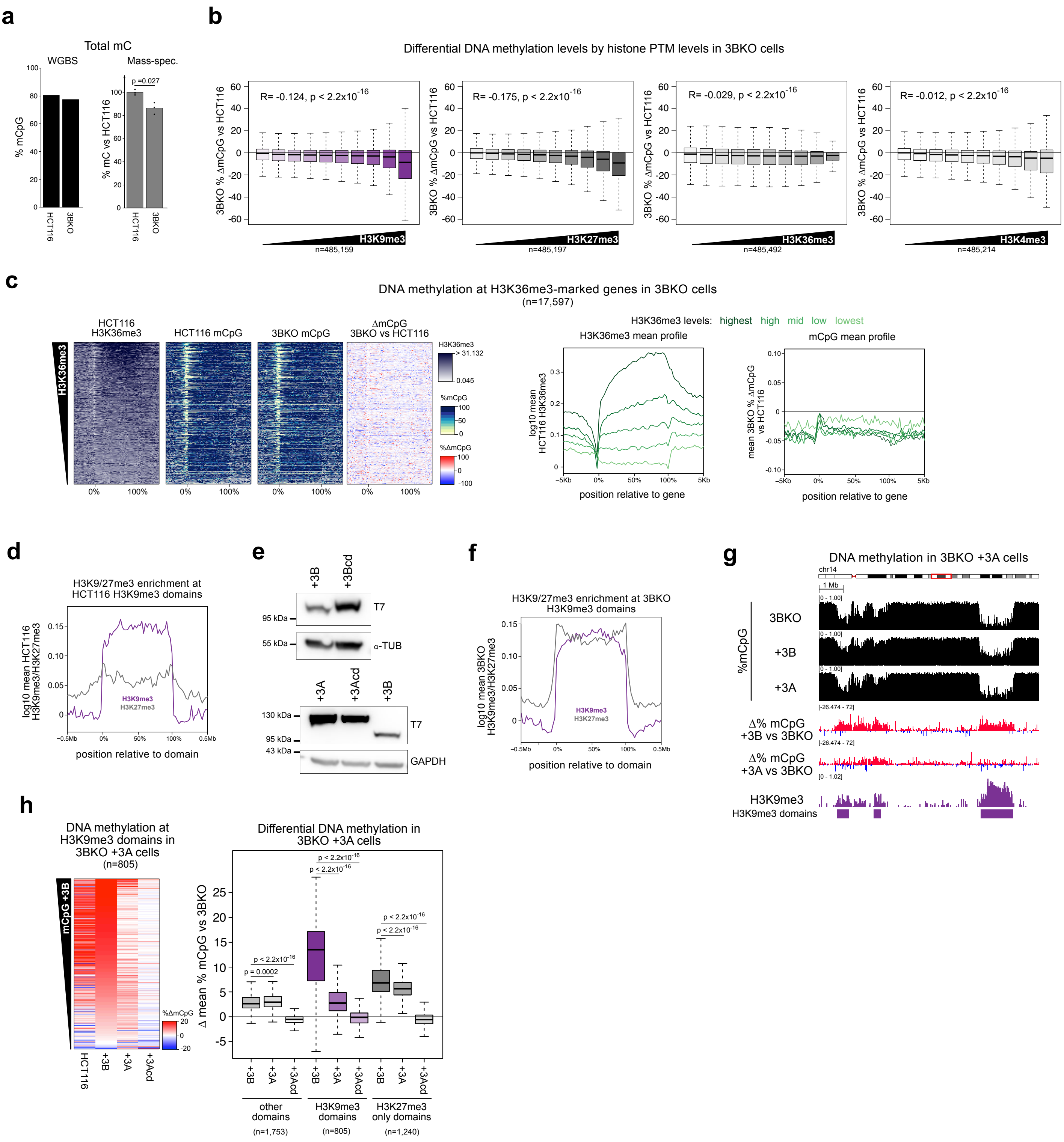

a
Total mC
WGBS
Mass-spec.
% mCpG
% mC vs HCT116
p =0.027
HCT116
3BKO
b
Differential DNA methylation levels by histone PTM levels in 3BKO cells
R= -0.124, p < 2.2x10-16
R= -0.175, p < 2.2x10-16
R= -0.029, p < 2.2x10-16
R= -0.012, p < 2.2x10-16
3BKO % ΔmCpG vs HCT116
H3K9me3
H3K27me3
H3K36me3
H3K4me3
n=485,159
n=485,197
n=485,492
n=485,214
c
DNA methylation at H3K36me3-marked genes in 3BKO cells
(n=17,597)
HCT116 H3K36me3
HCT116 mCpG
3BKO mCpG
ΔmCpG 3BKO vs HCT116
H3K36me3 levels: highest high mid low lowest
H3K36me3 mean profile
mCpG mean profile
log10 mean HCT116 H3K36me3
mean 3BKO % ΔmCpG vs HCT116
position relative to gene
d
H3K9/27me3 enrichment at HCT116 H3K9me3 domains
log10 mean HCT116 H3K9me3/H3K27me3
position relative to domain
e
+3B
+3Bcd
T7
α-TUB
95 kDa
55 kDa
+3A
+3Acd
GAPDH
130 kDa
43 kDa
f
H3K9/27me3 enrichment at 3BKO H3K9me3 domains
log10 mean 3BKO H3K9me3/H3K27me3
position relative to domain
g
DNA methylation in 3BKO +3A cells
chr14
1 Mb
%mCpG
3BKO
+3B
+3A
Δ% mCpG +3B vs 3BKO
Δ% mCpG +3A vs 3BKO
H3K9me3
H3K9me3 domains
h
DNA methylation at H3K9me3 domains in 3BKO +3A cells
(n=805)
mCpG +3B
HCT116
+3B
+3A
+3Acd
%ΔmCpG
Differential DNA methylation in 3BKO +3A cells
Δ mean % mCpG vs 3BKO
p = 0.0002
p < 2.2x10-16
other domains
(n=1,753)
H3K9me3 domains
(n=805)
H3K27me3 only domains
(n=1,240)

**Supplementary Figure 1**

**DNMT3B methylates heterochromatin**

**a)** Total DNA methylation levels are reduced in DNMT3B KO cells. Barplot of total methylated cytosine levels estimated by WGBS (left) and mass-spectrometry (right, mean methylation levels relative to HCT116 cells from 3 technical replicates). P- values are from two-sided T-tests. **b)** Correlation between loss of DNA methylation and enrichment of chromatin marks in DNMT3B KO cells. Boxplots showing difference in DNA methylation in DNMT3B KO to HCT116 cells at 2.5 kb genomic windows divided in deciles according to their histone modification enrichment calculated in HCT116 cells. The Pearson's correlation coefficient (R) is shown alongside its associated p-value. **c)** H3K36me3 enrichment at gene bodies does not correlate with DNA methylation loss in DNMT3B KO cells. Left, heatmaps showing levels of H3K36me3 and of absolute and differential DNA methylation at gene bodies in HCT116 and DNMT3B KO cells. Genes are ranked by their mean H3K36me3 levels. Right, profiles of H3K36me3 and differential DNA methylation levels at gene bodies, divided in 5 groups according to their H3K36me3 enrichment in HCT116 cells. **d)** Mean H3K9me3 and H3K27me3 ChIP-seq profiles at H3K9me3 domains in HCT116 cells. 82.9% of HCT116 H3K9me3 domains overlap with one or more H3K27me3 domains. **e)** Western blots showing ectopic expression of T7-tagged DNMT3B, DNMT3Bcd, DNMT3A and DNMT3Acd in DNMT3B KO cells. **f)** Mean H3K9me3 and H3K27me3 ChIP-seq profiles at H3K9me3 domains in DNMT3B KO cells. 92.6% of DNMT3B KO H3K9me3 domains overlap with one or more H3K27me3 domain. **g, h)** DNMT3B remethylates heterochromatin to a significantly higher level than DNMT3A. **g)** Representative genomic location showing gains of DNA methylation at H3K9me3 domains in DNMT3BKO cells expressing DNMT3B or DNMT3A. Genome browser plots show absolute (black) and differential (gain=red, loss=blue) DNA methylation levels, DNMT3B KO ChIP-seq signals and H3K9me3 domains defined in DNMT3B KO cells. ChIP-seq are normalised reads per $10^6$. **h)** Left, heatmaps of relative methylation levels at H3K9me3 domains. Values denote the change in methylation relative to DNMT3B KO cells. H3K9me3 domains are defined in DNMT3B KO cells and ranked by the mean gain of DNA methylation in DNMT3B KO cells expressing DNMT3B. Left, boxplots of DNA methylation difference to DNMT3B KO cells at H3K9me3, H3K27me3-only marked domains and the rest of the genome (other). +3B = DNMT3B KO + DNMT3B; +3A = DNMT3B KO + DNMT3A; +3Acd = DNMT3B KO + catalytically dead DNMT3A. For all the boxplots: lines = median; box = 25th–75th percentile; whiskers = 1.5 × interquartile range from box. P-values in **h.** are from two-sided Wilcoxon rank sum tests.

Fig. S2

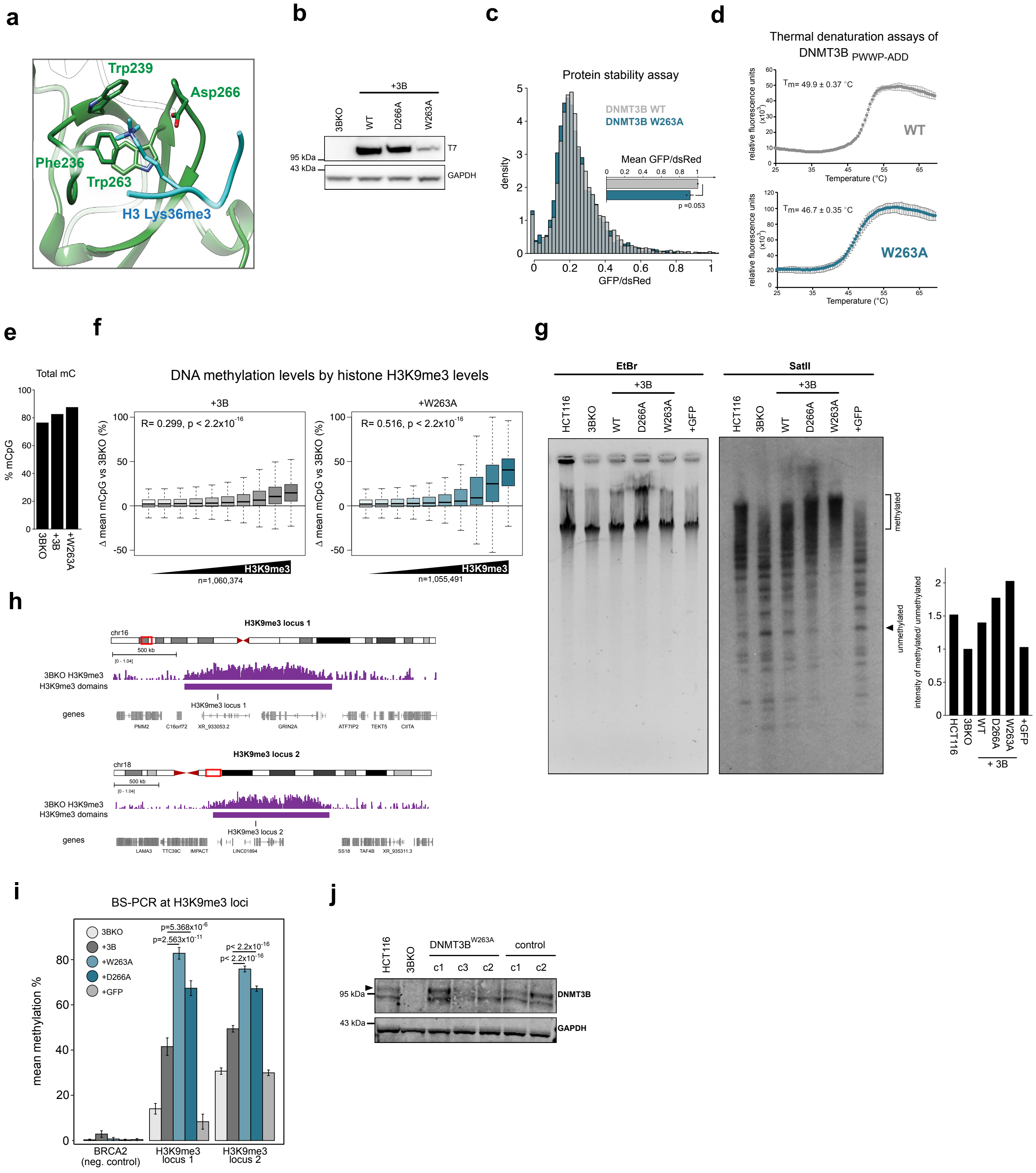
a
Trp239
Asp266
Phe236
Trp263
H3 Lys36me3
b
+3B
3BKO
WT
D266A
W263A
95 kDa
43 kDa
T7
GAPDH
c
Protein stability assay
DNMT3B WT
DNMT3B W263A
Mean GFP/dsRed
p =0.053
density
GFP/dsRed
d
Thermal denaturation assays of DNMT3B PWWP-ADD
Tm= 49.9 ± 0.37 °C
WT
Tm= 46.7 ± 0.35 °C
W263A
relative fluorescence units (x10³)
Temperature (°C)
e
Total mC
% mCpG
3BKO
+3B
+W263A
f
DNA methylation levels by histone H3K9me3 levels
+3B
R= 0.299, p < 2.2x10⁻¹⁶
+W263A
R= 0.516, p < 2.2x10⁻¹⁶
Δ mean mCpG vs 3BKO (%)
H3K9me3
n=1,060,374
n=1,055,491
g
EtBr
SatII
+3B
HCT116
3BKO
WT
D266A
W263A
+GFP
methylated
unmethylated
intensity of methylated/ unmethylated
+ 3B
h
H3K9me3 locus 1
chr16
500 kb
3BKO H3K9me3
H3K9me3 domains
genes
PMM2
C16orf72
XR_933053.2
GRIN2A
ATF7IP2
TEKT5
CIITA
H3K9me3 locus 2
chr18
LAMA3
TTC39C
IMPACT
LINC01894
SS18
TAF4B
XR_935311.3
i
BS-PCR at H3K9me3 loci
3BKO
+3B
+W263A
+D266A
+GFP
p=5.368x10⁻⁶
p=2.563x10⁻¹¹
p< 2.2x10⁻¹⁶
p< 2.2x10⁻¹⁶
mean methylation %
BRCA2 (neg. control)
H3K9me3 locus 1
H3K9me3 locus 2
j
HCT116
3BKO
DNMT3B W263A
control
c1
c3
c2
95 kDa
43 kDa
DNMT3B
GAPDH

**Supplementary Figure 2**

**Disruption of DNMT3B H3K36me3 binding causes gains of DNA methylation at heterochromatin**

**a)** Magnified view of DNMT3B ribbon representation of aromatic cage structure bound to trimethyl lysine 36 of H3 (blue) (Protein Data Bank code 5CIU). **b-d)** $DNMT3B^{W263A}$ is stable. **b)** Western blot of ectopically expressed T7-tagged DNMT3B proteins in DNMT3B KO cells. **c)** Stability of $DNMT3B^{WT}$ and $DNMT3B^{W263A}$ measured by fluorescence reporter. Histogram shows the density distribution of single cell GFP/dsRed ratios of one representative experiment. The barplot shows the mean GFP/dsRed ratios of all the cells, measurements for the mutant normalized to the wild-type, for three independent experiments. P value is from Student's T-test. **d)** Thermal denaturation assays on purified $DNMT3B^{WT}$ or $DNMT3B^{W263A}$ protein PWWP and ADD fragments. Graphs showing SYBR-safe fluorescence measured at temperatures from 25°C to 69.5°C. Mean and standard deviation of three experiments are plotted with mean Tm values stated. $DNMT3B^{WT}$ data are repeated from Fig 4e as the data were part of the same experiment. **e)** Total levels of DNA methylation measured by WGBS analysis. +3B = $DNMT3B^{WT}$ cells; +W263A = $DNMT3B^{W263A}$ cells. **f)** Boxplots showing gains of DNA methylation in $DNMT3B^{WT}$ or $DNMT3B^{W263A}$ cells at 2.5 kb genomic windows ranked according to H3K9me3 enrichment in DNMT3B KO cells before being grouped into deciles. Lines = median; box = 25th–75th percentile; whiskers = 1.5 × interquartile range from box. Pearson's correlations and associated p-values are shown. **g-i)** Expression of $DNMT3B^{D266A}$ leads to hypermethylation of heterochromatin. **g)** Methylation sensitive Southern blot showing reduced digestions of satellite II sequences in $DNMT3B^{W263A}$ or $DNMT3B^{D266A}$ cells compared to $DNMT3B^{WT}$, DNMT3B KO cells and DNMT3B KO cells expressing GFP. Ethidium bromide stained gel (EtBr) is shown as loading control. Barplot shows signal quantification of satellite II Southern blot using the ratio of the methylated over unmethylated regions indicated. **h)** Genomic location of the two non-repetitive H3K9me3 loci assayed by BS-PCR in this study. H3K9me3 ChIP-seq signals in DNMT3B KO cells are shown above the amplicon locations. **i)** Mean methylation by BS-PCR at H3K9me3 loci alongside the H3K4me3-marked BRCA2 promoter in DNMT3B KO cells expressing DNMT3B mutants. P-values are from two-sided Wilcoxon rank sum tests. The number of reads analysed per each sample are shown in *Supplementary Table 4*. **j)** Western blot showing DNMT3B expression in CRISPR/Cas9 edited clones. Both DNMT3B2 (black arrow) and the catalytically inactive DNMT3B3 are visible.

### Fig. S3

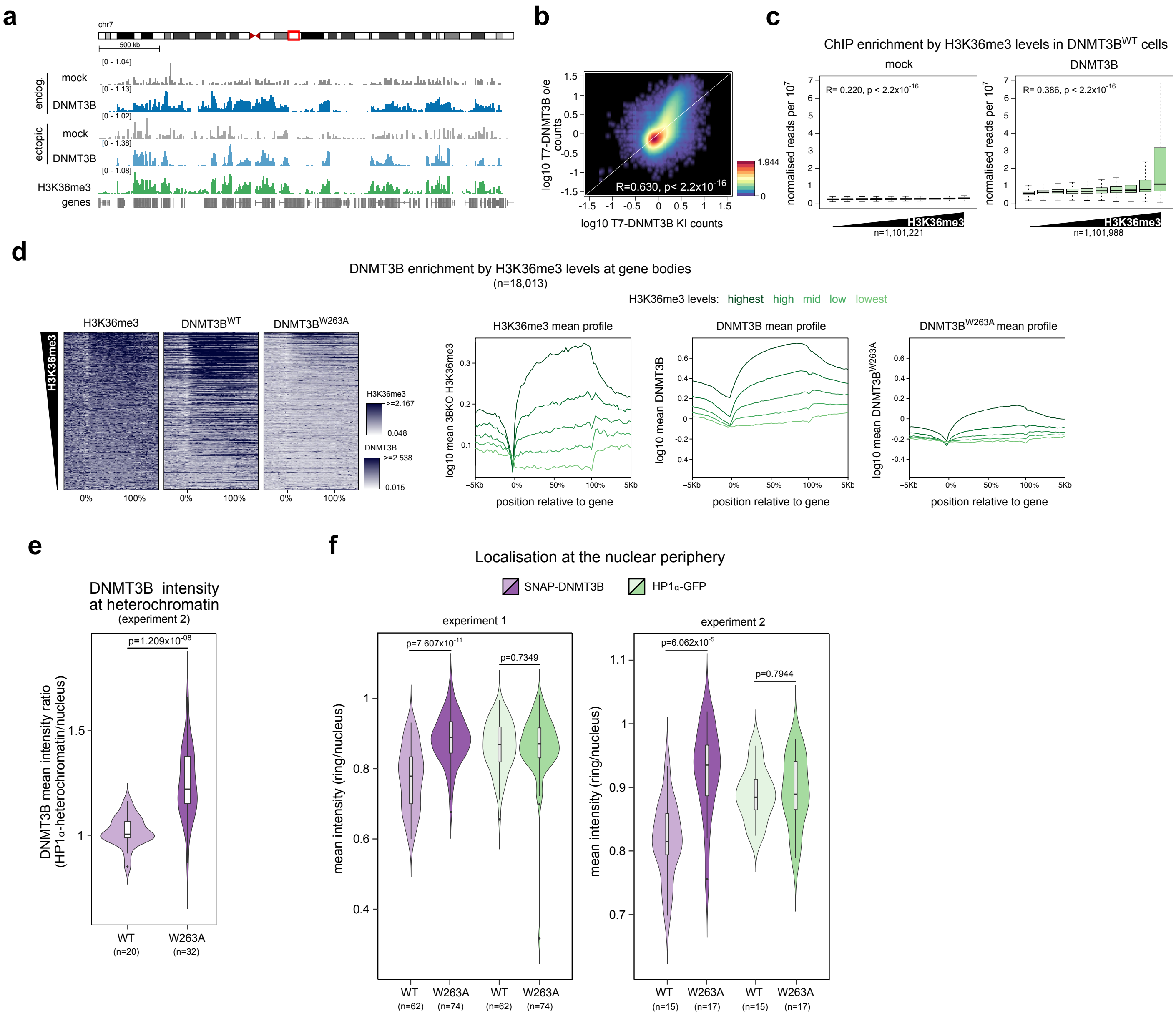
a
chr7
500 kb
endog.
mock
DNMT3B
ectopic
mock
DNMT3B
H3K36me3
genes
b
log10 T7-DNMT3B o/e counts
log10 T7-DNMT3B KI counts
R=0.630, p< 2.2x10-16
c
ChIP enrichment by H3K36me3 levels in DNMT3BWT cells
mock
DNMT3B
R= 0.220, p < 2.2x10-16
R= 0.386, p < 2.2x10-16
normalised reads per 107
H3K36me3
n=1,101,221
n=1,101,988
d
DNMT3B enrichment by H3K36me3 levels at gene bodies
(n=18,013)
H3K36me3 levels: highest high mid low lowest
H3K36me3
DNMT3BWT
DNMT3BW263A
H3K36me3 mean profile
DNMT3B mean profile
DNMT3BW263A mean profile
log10 mean 3BKO H3K36me3
log10 mean DNMT3B
log10 mean DNMT3BW263A
position relative to gene
e
DNMT3B intensity at heterochromatin
(experiment 2)
p=1.209x10-08
DNMT3B mean intensity ratio (HP1α-heterochromatin/nucleus)
WT (n=20)
W263A (n=32)
f
Localisation at the nuclear periphery
SNAP-DNMT3B
HP1α-GFP
experiment 1
experiment 2
p=7.607x10-11
p=0.7349
p=6.062x10-5
p=0.7944
mean intensity (ring/nucleus)
WT (n=62)
W263A (n=74)
WT (n=62)
W263A (n=74)
WT (n=15)
W263A (n=17)
WT (n=15)
W263A (n=17)

**Supplementary Figure 3**

**Impaired H3K36me3 binding leads to increased DNMT3B localisation to heterochromatin**

**a-c)** Ectopically expressed DNMT3B recapitulates the genome binding profile of endogenous DNMT3B. **a)** Binding profiles of endogenous T7-DNMT3B, ectopic T7-DNMT3B and respective mock at a representative genomic region. Genome browser plots showing T7-DNMT3B ChIP signals along with H3K36me3 ChIP-seq in HCT116 cells, normalised reads per $10^6$. **b)** Density scatter plot showing genome-wide correlations in 2.5 kb windows between endogenous T7-DNMT3B and T7-DNMT3B ectopically expressed in HCT116 cells. Correlation was calculated based on log10 transformed normalised read counts. Pearson's correlation (R) and associated p-value is shown. **c)** Boxplot showing mock or T7-DNMT3B$^{WT}$ levels normalised over input at 2.5 kb genomic windows of increased H3K36me3 enrichment in DNMT3B KO cells before being grouped into deciles. Lines = median; box = 25th–75th percentile; whiskers = 1.5 × interquartile range from box. Pearson's correlation and associated p-value is shown. **d)** Decreased enrichment of DNMT3B$^{W263A}$ at H3K36me3 marked gene bodies. Left, heatmaps showing the levels of H3K36me3 in DNMT3B KO cells, T7-DNMT3B$^{WT}$ or T7-DNMT3B$^{W263A}$ at gene bodies. Genes are ranked by their mean H3K36me3 levels. Right, mean profiles of H3K36me3 or T7-DNMT3B proteins levels at gene bodies, divided in 5 groups according to their H3K36me3 enrichment. **e)** Violin plot showing the distribution of DNMT3B mean intensity ratio between HP1α-marked heterochromatin and the rest of the nucleus from the second replicate experiment. Lines = median; box = 25th–75th percentile; whiskers = 1.5 × interquartile range from box. P-value is from two-sided Wilcoxon rank sum test. **f)** DNMT3B localisation at the nuclear periphery. Violin plot showing the intensity of DNMT3B or HP1α in the 0.5 μm outermost nuclear ring as proportion of the total nuclear intensity. Two independent replicate experiments are shown. Lines = median; box = 25th–75th percentile; whiskers = 1.5 × interquartile range from box. P-values are from two-sided Wilcoxon rank sum tests.

### Fig. S4

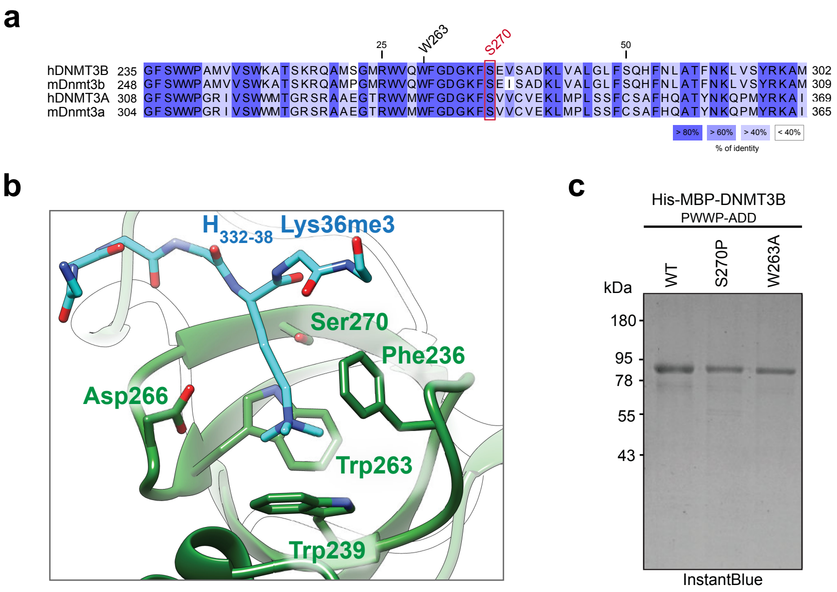
a
25
W263
S270
50
hDNMT3B 235 GFSWWPAMVVSWKATSKRQAMSGMRWVQWFGDGKFSEVSADKLVALGLFSQHFNLATFNKLVSYRKAM 302
mDnmt3b 248 GFSWWPAMVVSWKATSKRQAMPGMRWVQWFGDGKFSEISADKLVALGLFSQHFNLATFNKLVSYRKAM 309
hDNMT3A 308 GFSWWPGRIVSWWMTGRSRAAEGTRWVMWFGDGKFSVVCVEKLMPLSSFCSAFHQATYNKQPMYRKAI 369
mDnmt3a 304 GFSWWPGRIVSWWMTGRSRAAEGTRWVMWFGDGKFSVVCVEKLMPLSSFCSAFHQATYNKQPMYRKAI 365
> 80%
> 60%
> 40%
< 40%
% of identity
b
H332-38 Lys36me3
Ser270
Phe236
Asp266
Trp263
Trp239
c
His-MBP-DNMT3B
PWWP-ADD
WT
S270P
W263A
kDa
180
95
78
55
43
InstantBlue

**Supplementary Figure 4**

**The ICF1 syndrome S270P mutation destabilises DNMT3B**

**a)** Multiple sequence alignment of part of the DNMT3B PWWP domain protein sequence with DNMT3A and mouse orthologues showing the location of serine 270. **b)** Magnified view of DNMT3B histone H3 binding pocket with the ICF1-mutated Ser-270 indicated and H3 Lys 36 shown in blue. **c)** SDS-PAGE gel of the proteins used in Fig. 4e stained with InstantBlue protein stain.

### Fig. S5

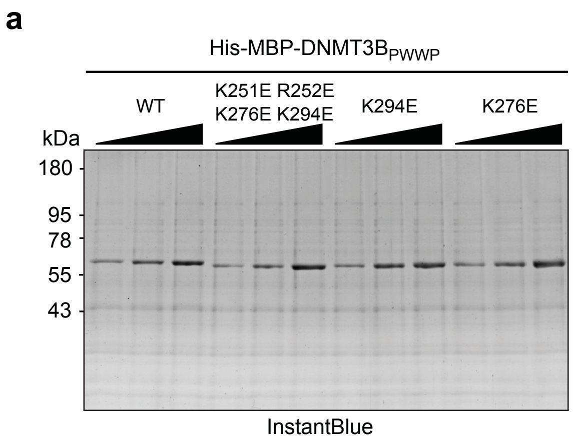
a
His-MBP-DNMT3B$_{PWWP}$
WT
K251E R252E
K276E K294E
K294E
K276E
kDa
180
95
78
55
43
InstantBlue

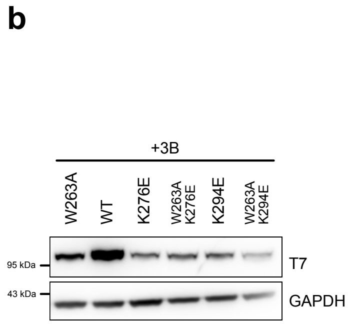
b
+3B
W263A
WT
K276E
W263A
K276E
K294E
W263A
K294E
95 kDa
43 kDa
T7
GAPDH

**Supplementary Figure 5**

**DNMT3B-PWWP binding to DNA is dispensable for localisation to heterochromatin**

**a)** SDS-PAGE gel stained with InstantBlue protein stain of proteins used in Fig. 5. **b)** Western blot of ectopically expressed T7-tagged DNMT3B proteins in DNMT3B KO cells.

### Fig. S6

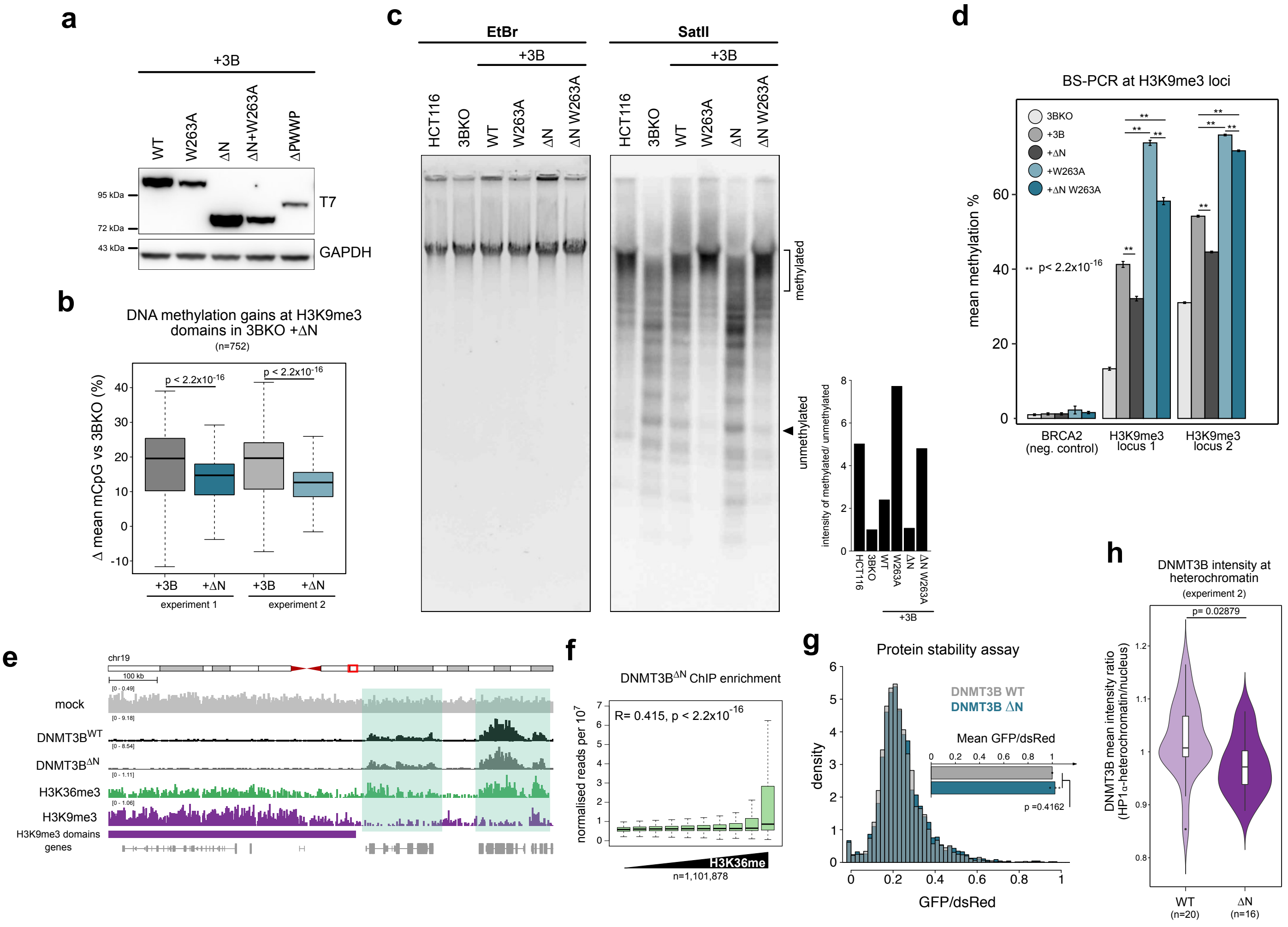
a
+3B
WT
W263A
ΔN
ΔN+W263A
ΔPWWP
95 kDa
72 kDa
43 kDa
T7
GAPDH
b
DNA methylation gains at H3K9me3 domains in 3BKO +ΔN
(n=752)
Δ mean mCpG vs 3BKO (%)
p < 2.2x10-16
p < 2.2x10-16
+3B
+ΔN
+3B
+ΔN
experiment 1
experiment 2
c
EtBr
SatII
+3B
+3B
HCT116
3BKO
WT
W263A
ΔN
ΔN W263A
methylated
unmethylated
intensity of methylated/ unmethylated
d
BS-PCR at H3K9me3 loci
3BKO
+3B
+ΔN
+W263A
+ΔN W263A
** p< 2.2x10-16
mean methylation %
BRCA2
(neg. control)
H3K9me3
locus 1
H3K9me3
locus 2
e
chr19
100 kb
mock
DNMT3BWT
DNMT3BΔN
H3K36me3
H3K9me3
H3K9me3 domains
genes
f
DNMT3BΔN ChIP enrichment
R= 0.415, p < 2.2x10-16
normalised reads per 10^7
H3K36me
n=1,101,878
g
Protein stability assay
DNMT3B WT
DNMT3B ΔN
Mean GFP/dsRed
p =0.4162
density
GFP/dsRed
h
DNMT3B intensity at heterochromatin
(experiment 2)
p= 0.02879
DNMT3B mean intensity ratio
(HP1α-heterochromatin/nucleus)
WT
(n=20)
ΔN
(n=16)

**Supplementary Figure 6**

**The N-terminus facilitates methylation of heterochromatin by DNMT3B**

**a)** Western blot of ectopically expressed T7-tagged DNMT3B proteins in DNMT3B KO cells. **b)** Boxplot showing gains of DNA methylation at H3K9me3 domains in DNMT3B$^{WT}$ or DNMT3B$^{\Delta N}$ cells from two independent experiments. Lines = median; box = 25$^{th}$–75$^{th}$ percentile; whiskers = 1.5 × interquartile range from box. P-values are from two-sided Wilcoxon rank sum tests. **c)** Methylation sensitive Southern blot showing digestion of satellite II sequences in DNMT3B KO cells expressing DNMT3B mutants (centre). Ethidium bromide stained gel (EtBr) is shown as a loading control (left). Barplot shows signal quantification of satellite II Southern blot using the ratio of the methylated over unmethylated regions indicated (right). **d)** Mean methylation by BS-PCR at H3K9me3 loci alongside the H3K4me3-marked BRCA2 promoter in DNMT3B mutant cells. P-values are from two-sided Wilcoxon rank sum tests. The number of reads analysed per each sample are shown in *Supplementary Table 4*. **e)** Genome browser plots showing DNMT3B$^{WT}$ and DNMT3B$^{\Delta N}$ ChIP signal and H3K36me3 and H3K9me3 ChIP-seq signal from DNMT3B KO cells, normalised reads per $10^6$. Green rectangles highlight similar enrichment profiles for DNMT3B$^{WT}$ and DNMT3B$^{\Delta N}$ at H3K36me3-rich loci. **f)** Boxplot showing levels of T7-DNMT3B$^{\Delta N}$ normalised over input at 2.5 kb genomic windows of increased H3K36me3 enrichment and grouped into deciles. Lines = median; box = 25th–75th percentile; whiskers = 1.5 × interquartile range from box. Pearson's correlation is shown. **g)** Stability of DNMT3B$^{WT}$ and DNMT3B$^{\Delta N}$ measured by fluorescence reporter. Histogram shows the density distribution of single cell GFP/dsRed ratios of one representative experiment. The barplot shows the mean GFP/dsRed ratios of all the cells, measurements for the mutant normalized to the wild-type, for three independent experiments. P value is from Student's T-test. **h)** Violin plot showing the distribution of DNMT3B mean intensity ratio between HP1α-marked heterochromatin and the rest of the nucleus from the second replicate experiment. For boxplots, lines = median; box = 25th–75th percentile; whiskers = 1.5 × interquartile range from box and p-value are from two-sided Wilcoxon rank sum tests. DNMT3B$^{WT}$ data are repeated from *Fig. S3e* as the data are part of the same experiment.
